# Exploring the role of cultivar, year and plot age in the incidence of esca and Eutypa dieback: insights from 20 years of regional surveys in France

**DOI:** 10.1101/2024.03.19.585220

**Authors:** Lucas Etienne, Frédéric Fabre, Davide Martinetti, Elise Frank, Lucie Michel, Valérie Bonnardot, Lucia Guérin-Dubrana, Chloé E. L. Delmas

**Affiliations:** INRAE, ISVV, Bordeaux Sciences Agro, Santé et Agroécologie du Vignoble, 33140 Villenave d’Ornon, France; INRAE, Biostatistiques et Processus Spatiaux, 84000 Avignon, France; Plateforme ESV, INRAE, Biostatistiques et Processus Spatiaux, 84914 Avignon, France; CNRS, Université Rennes 2, Littoral Environnement Télédétection Géomatique, 35045 Rennes, France

**Keywords:** esca, Eutypa dieback, epidemiology, INLA, surveillance, *Vitis vinifera*

## Abstract

Grapevine trunk diseases cause yield losses and vine mortality in vineyards worldwide. However, there have been few quantitative studies evaluating grapevine dieback on a large spatial and temporal scale. Here, we consolidated and standardized databases from the 13 main wine regions of France, compiling records of leaf symptoms associated with esca and Eutypa dieback from 2082 plots and 36 cultivars over a 20-year period. This large dataset was used (a) for quantitative analysis of the prevalence (number of plots with at least one symptomatic plant) and incidence (percentage of symptomatic plants) of esca and Eutypa dieback; and (b) to decipher the effects of cultivar, year and plot age on both the prevalence and incidence of esca leaf symptoms by temporal Bayesian modelling. Esca was present on a mean of 74 ± 2% plots annually, with an incidence of 3.1 ± 0.1%. Eutypa dieback occurred in 41 ± 3% of the plots, with an incidence of 1.4 ± 0.1%. Our modelling approach revealed that the cultivar had a significant impact on the prevalence of esca, but not on its incidence when prevalence is greater than zero. Esca prevalence remained stable, whereas esca incidence was higher than the mean value in six of the years after 2012. We also found a significant non-linear effect of plot age, with 10- to 30-year-old plots significantly more susceptible, depending on the cultivar. This study clearly illustrates the importance of considering extensive and continuous monitoring to improve our understanding of the impact and evolution of crop diseases.

## I) Introduction

Perennial plant dieback is characterised by the premature and progressive death of shoots, loss of plant vigor, and, ultimately, plant death. Abiotic factors, such as drought, have been shown to be a major cause of dieback in perennial plants (Allen et al., 2010; Cailleret et al., 2019; Hammond et al., 2022), as well as pathogens (Ciesla and Donaubauer, 1994) and their interactions (Jurskis, 2005). Dieback is a well-known phenomenon in forest ecology, observed in various regions of the world (Allen, 2009; Hammond et al., 2022; Hartmann et al., 2018). However, there have been no large-scale quantitative studies assessing dieback in perennial crops.

Dieback of grapevine (*Vitis vinifera* L.) is caused by various abiotic and biotic factors, including diseases affecting grapevine wood. Trunk diseases, such as esca, Eutypa dieback, and Botryosphaeriaceae dieback, are the most widespread globally (Guerin-Dubrana et al., 2019; Mugnai et al., 1999). These diseases cause vine destruction leading to yield losses (Bertsch et al., 2009; Gramaje et al., 2018; Mondello et al., 2018), although these can be overestimated in some regions (Dewasme et al., 2022). The current epidemic spread of trunk diseases in Europe can be traced back to the early 1990s (Mugnai et al., 1999; Reisenzein et al., 2000; Surico et al., 2000). In the first few years of the 21st century, until at least 2008, the incidence of grapevine trunk diseases, including esca in particular, appears to have increased (Bruez et al., 2013). This increase is a matter of great concern to vine growers, particularly in the wake of the sodium arsenate ban introduced in France in 2001, as this was the only effective treatment for esca (Mugnai et al., 1999). Several factors may be associated with variation in the incidence of trunk diseases. These factors include cultivar (Gastou et al., 2024), year (Dewasme et al., 2022) and plot age (Fussler et al., 2008). Cultivar is a major factor underlying differences in the incidence of esca disease between vineyards and vine-growing regions. Indeed, there was a considerable variability of the proportion of symptomatic plants per cultivar in France during the monitoring of 46 cultivars planted in a common garden vineyard over a period of seven years (Gastou et al., 2024), in Italy during the monitoring of 67 cultivars for one year (Murolo and Romanazzi, 2014) and in Spain, during the monitoring of 47 cultivars for three years (Chacón-Vozmediano et al., 2021). Certain cultivars, such as Merlot, only rarely display esca leaf symptoms whereas others, such as Sauvignon Blanc, are frequently affected. Moreover, strong interannual variability has been observed within vineyards (Calzarano et al., 2018; Dewasme et al., 2022). The age of the vines significantly influences their response to biotic and abiotic stresses, determining their tolerance or susceptibility. As a result, the incidence and expression of symptoms of stress increase linearly with plant age (Pandey et al., 2015). The effect of vineyard age on esca incidence is unclear. A few studies have reported an absence of correlation between plot age and esca incidence (Bruez et al., 2013; Péros et al., 2008), but others have reported a significant age effect for plots aged from 10 to 21 years (Kovács et al., 2017). By contrast, other studies have suggested that the relationship between plot age and symptom expression is quadratic rather than linear, with disease incidence highest at intermediate ages (Fulchin et al., 2019; Fussler et al., 2008). Incidence appears to be higher in vineyards of between 15 and 25 years of age than in vineyards of other ages, as shown in a study of 22 cultivars (Fussler et al., 2008) and, more recently, in a study of five cultivars (Fulchin et al., 2019). However, in these studies, age was considered as a categorical ordered variable (e.g. ‘young’ if the plot was less than 7 years old and ‘old’ if more than 11 years old, Romanazzi et al., 2009, or 0-15, 15-25, 25-40, and over 40 years old, Fussler et al., 2008). No study has ever addressed age as a fully continuous quantitative variable, probably due to a lack of long-term, large-scale monitoring.

Distinguishing between the effects of year and plot age requires large-scale and long-term monitoring, to ensure that the variables are not correlated, or at least no more than weakly correlated. Large-scale monitoring is essential to improve pest surveillance for plants, particularly over large spatial scales, and to facilitate the implementation of effective measures for preventing the spread of pathogens and insect pests and for controlling epidemics (Carvajal-Yepes et al., 2019; Mariette et al., 2023; Parnell et al., 2017). General surveillance encompasses the collection and analysis of information on plant disease and plays a crucial role in the detection and effective management of pathogens (Aguayo et al., 2021; ISPM 6 (FAO) 1997; Parnell et al., 2017). The extensive monitoring of significant pathogens, including native ones, is also crucial for obtaining spatial indicators of vineyard health, and for tracking temporal trends. This information enables managers and policymakers to implement sustainable management practices in vineyards. In France, a National Grapevine Trunk Diseases Survey was performed between 2003 and 2008, to monitor grapevine trunk disease incidence and mortality and to assess its significance for viticulture in seven vine-growing regions This survey included 12 cultivars and 329 vineyard plots (Bruez et al., 2013; Fussler et al., 2008). However, the collection of more recent information over a longer period would be required to assess progression of grapevine trunk disease levels over the last few years.

In this study, we collected and curated different databases from the 13 main wine regions of France, to generate a unified database for leaf symptoms of two major trunk diseases, esca and Eutypa dieback, covering a period of 20 years (2003 to 2022). This unified national database covers 2082 plots and 36 cultivars and was used to describe the prevalence (percentage of plots with at least one symptomatic plant) and incidence (percentage of symptomatic plants per plot), as defined by Nutter et al. (2006), of esca and Eutypa dieback over the different years, cultivars, regions, and vineyard ages. Furthermore, the time-series data collected at plot scale were subjected to modelling by the integrated nested Laplace approximation (INLA) method (Rue et al., 2009, 2017) to explore the effects of cultivar, year and plot age on both the prevalence and incidence of esca whilst accounting for temporal dependencies.

## II) Materials and methods

### 1) French database of grapevine trunk disease observations

The database contains 12,587 observations of leaf symptoms of esca and Eutypa dieback, in the main wine-producing regions in France. This database is stored in the information system of the French Epidemiological Plant Health Surveillance Platform (ESV Platform). The observations were obtained from two sources: (i) regional surveys conducted from 2003 to 2022, and (ii) the historical “National Grapevine Trunk Diseases Survey”, which tracked the progression of grapevine trunk diseases throughout France from 2003 to 2008 (Fussler et al., 2008; Grosman and Doublet, 2012; Bruez et al., 2013). The historical and regional surveys were conducted by experts from diverse public or private agronomic institutes or associations in each of the regions.

Observations of esca leaf symptoms were obtained in 884 different municipalities (2.4 ± 2.5 plots per municipality, mean ± standard deviation, SD), in 49 provinces, 13 vine-growing regions, and 10 administrative regions (see Figure 1 and Table S1 for the number of plots per region and monitoring years). They took place at the end of August, which corresponds to the period of maximum cumulative incidence in French vineyards (see the intra-seasonal dynamics presented in Lecomte et al. 2024). Plants were scored symptomatic when typical leaf stripe symptoms were observed as presented in Lecomte et al. (2024). Thirty-six different cultivars were monitored, with a mean of 58 ± 69 plots per cultivar, as described in Table S2. Three cultivars (Italia, Alphonse Lavallée and Sauvignon Gris) were monitored on only one plot each and were therefore excluded from the analysis. Moreover, only two plots were monitored in the Vendée wine region (Chenin cultivar), and these plots were also, therefore, excluded from the analysis. Finally, mean plot age was 27 ± 13 years, and plot age ranged from 1 to 101 years (Figure 2). However, the date of plantation was not recorded for 25% of the plots. Each region was represented by a different set of cultivars (Figure S1).

**Figure 1:**
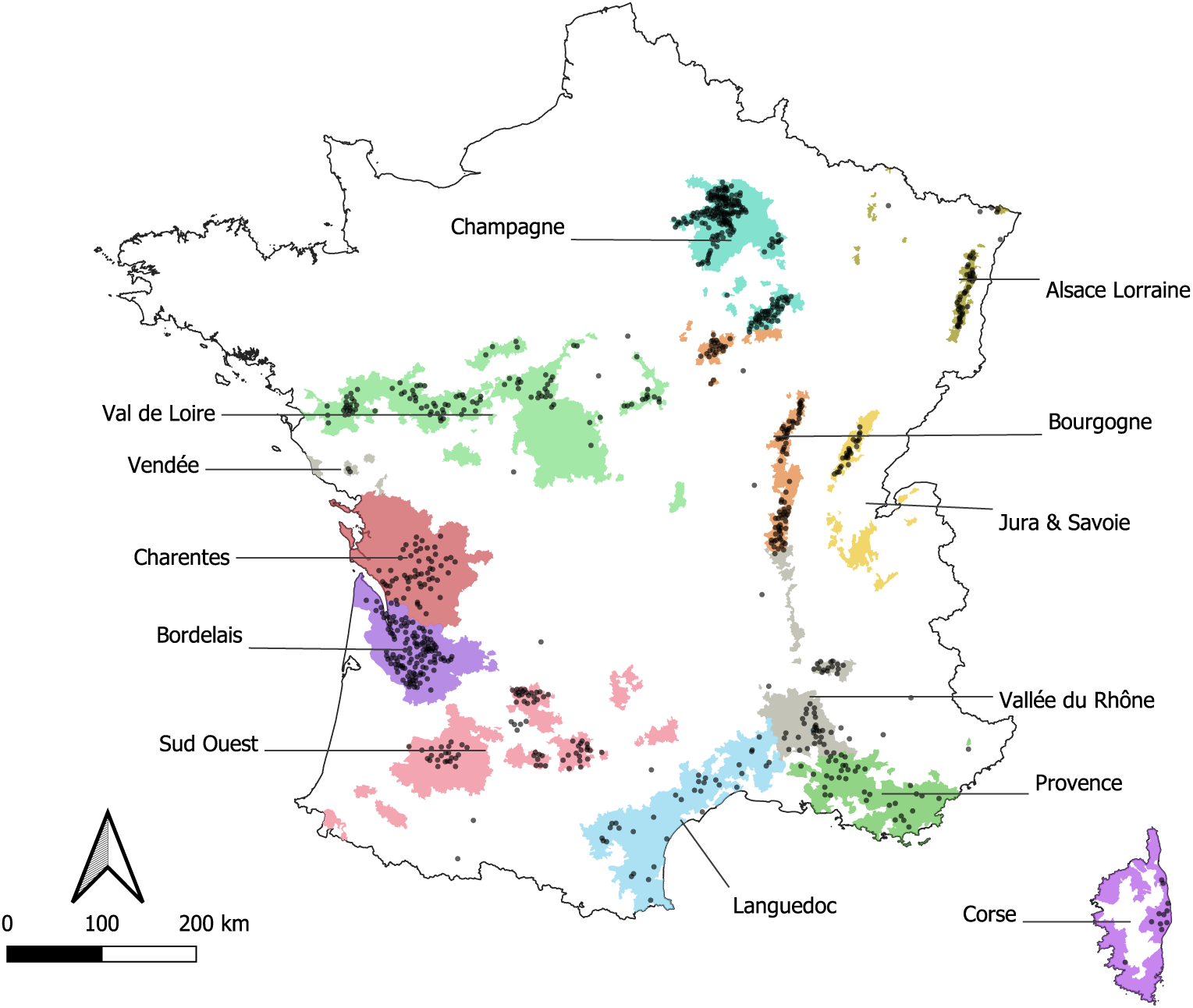
Map of the French wine-growing regions showing the locations of the 884 municipalities in which plots were monitored for esca leaf symptoms (black dots). These municipalities are located in 49 provinces, 13 vine-growing regions, and 10 administrative regions. Eutypa dieback was also monitored in 592 of these municipalities in 39 provinces, 10 wine regions, and 7 administrative regions.

**Figure 2:**
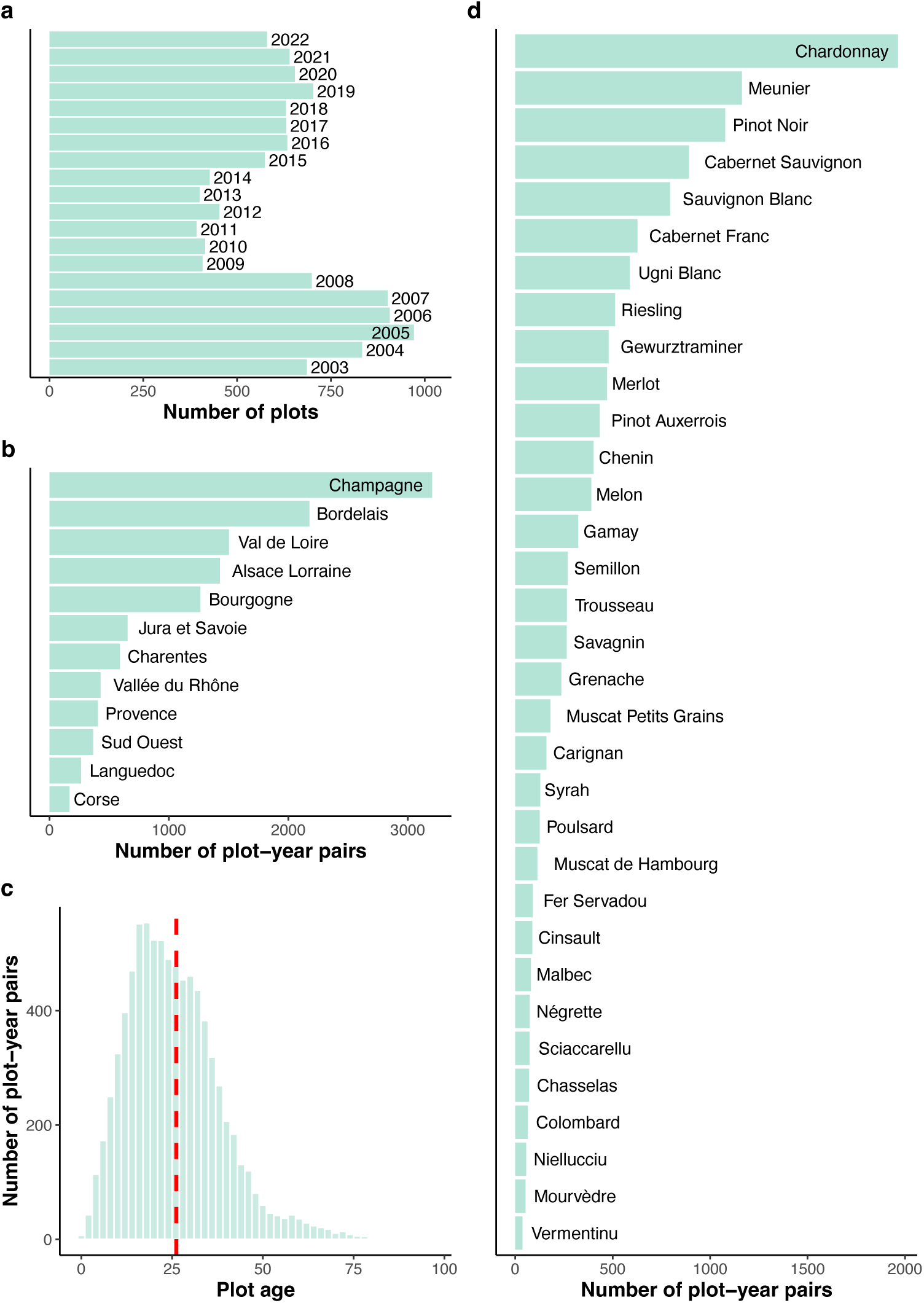
Number of plots or observations (plot-year pairs) monitored in France for esca between 2003 and 2022 and recorded in the database, by year (a), wine region (b), plot age (c), and *Vitis vinifera* cultivar (d). An observation corresponds to one plot monitored in a given year. The red dotted line in (c) represents the mean age.

For 63% of the plots monitored for esca, annotation was also available for Eutypa dieback symptoms. Eutypa dieback was monitored between April and May (Table S1) to score plants presenting stunted shoots with necrotic leaves as shown in Mondello et al. (2018). In total 7,073 observations were available for this disease, spread over 1,310 plots located in 592 municipalities (2.2 ± 2.4 (mean ± SD) plots per municipality), 39 provinces, 10 wine regions, and 7 administrative regions. All cultivars other than Meunier, Sciaccarellu, Vermentinu, and Niellucciu were monitored for Eutypa dieback, with data available only for esca leaf symptoms for these exceptions.

Mortality was recorded, but the methods used to score dead plants were not homogeneous across regions. We, therefore, discarded these data from the analysis. Similarly, information on rootstock, pruning technique, young replanted vines, and apoplexy (total dehydration of the canopy) was scarce and inconsistently reported, and such data were not, therefore, considered in this analysis.

The number of vines with esca or Eutypa dieback symptoms on leaves was recorded on a defined number of vines in the plots described above. A mean of 420 ± 530 vines was observed per plot (see supplementary materials, Table S2). One plot of Cabernet Sauvignon was monitored in its entirety as part of an experiment, accounting for the large mean number of vines observed for this cultivar (Table S2). The definitions of disease prevalence and incidence proposed by Nutter et al. (2006) were used.

### 2) Estimations of the prevalence and incidence of leaf symptoms of esca and Eutypa dieback

Disease prevalence was calculated, for a given year, as the percentage of plots on which esca, or Eutypa dieback (depending on the disease considered for the analysis) was observed on at least one plant. Disease incidence was calculated for each plot in a given year by dividing the number of plants presenting esca leaf symptoms (as described by Lecomte et al. 2012) or Eutypa dieback leaf symptoms (as described by Sosnowski et al. 2007) by the number of plants monitored in the plot concerned (included dead or missing vines).

### 3) Deciphering the effects of cultivar, year, and plot age on the prevalence and incidence of esca leaf symptoms

In addition to the descriptive analysis of the whole database, a statistical model was developed to estimate the effect of cultivar, year and plot age on both prevalence and incidence of esca leaf symptoms at plot scale. In this modelling approach, only esca was used, as Eutypa dieback is affecting almost exclusively the Ugni Blanc cultivar and presents a decreasing incidence over time, in contrast to esca (Figure 3). To optimise model identifiability, we applied some criteria to select the data used in this approach. We selected observations (one plot in a given year) from plots under 50 years old (older plots excluded), for which at least five years of observation were available (not necessarily consecutive), and cultivars for which at least 300 observations were available. The application of these criteria resulted in a database of 5161 observations (from the 12587 observations initially identified) spread over 12 cultivars (Table 1). On average, a plot was monitored for 9 ± 4 [5, 20] years. Mean plot age ranged from 18 to 32 years (Table 1). Mean plot age tended to increase from 2003 to 2013, remaining stable thereafter (Figure 4).

**Figure 3:**
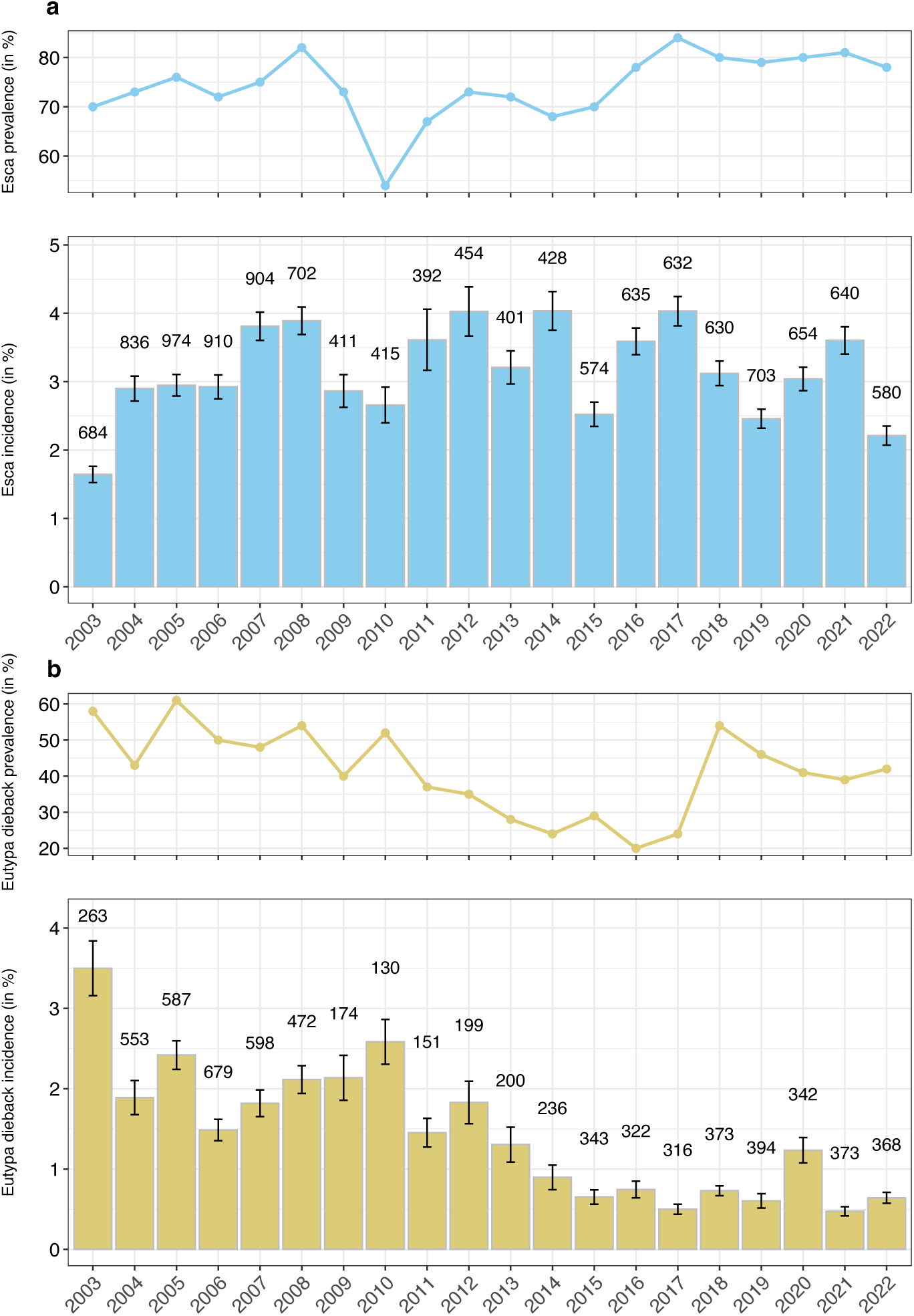
The mean prevalence and incidence of esca and Eutypa dieback leaf symptoms per year in France over the 2003-2022 period. The prevalence (shown as curves in the top panel) was determined as the number of plots with at least one symptomatic vine recorded per year. The incidence (represented by bar charts in the bottom panel) was determined as the percentage of symptomatic vines observed per plot each year. The numbers displayed correspond to the number of plots monitored for the incidence of (a) esca (blue dots, lines and bars) and (b) Eutypa dieback (mustard dots, lines and bars). The error bars indicate the standard error of the mean.

**Figure 4:**
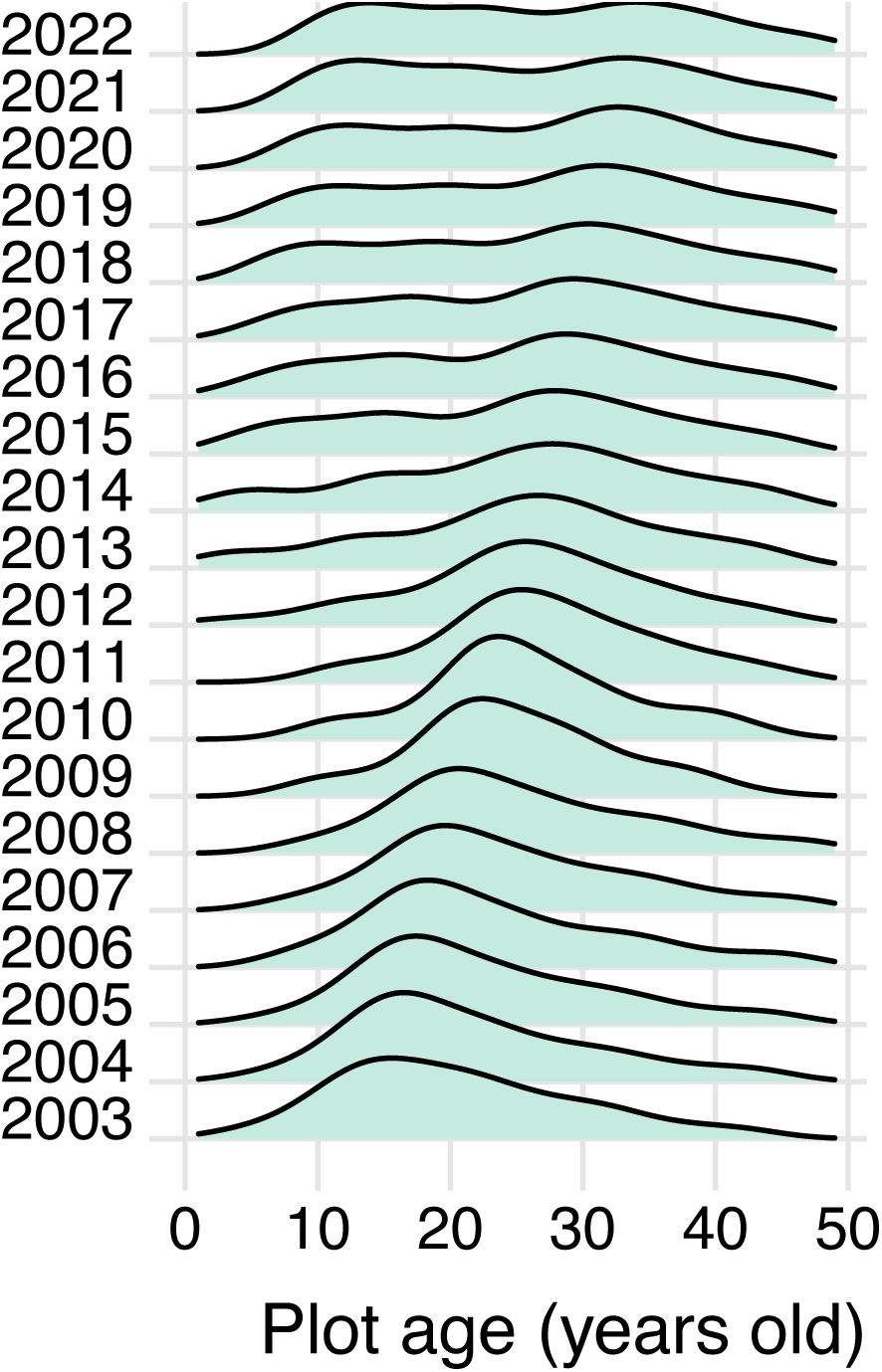
Distribution of plot age (range: 1 to 49 years) for each year for the 12 *Vitis vinifera* cultivars used in the modelling approach.

**Table 1:** Summary statistics for plot age and number of plots (total and per year) monitored for the 12 *Vitis vinifera* cultivars retained for statistical modelling. Maximum plot age was fixed at 49 years.

| Cultivar | Plot age (mean ± SD [min, max] years) | Total number of plots | Mean (± SD) number of plots per year |
| --- | --- | --- | --- |
| Cabernet Sauvignon | 23 ± 9 [2, 49] | 85 | 14 ± 13 |
| Sauvignon Blanc | 20 ± 10 [3, 49] | 62 | 8 ± 10 |
| Chardonnay | 30 ± 9 [7, 49] | 61 | 6 ± 10 |
| Cabernet Franc | 23 ± 13 [2, 49] | 53 | 9 ± 12 |
| Merlot | 24 ± 10 [7, 49] | 52 | 13 ± 11 |
| Pinot Noir | 29 ± 9 [7, 49] | 47 | 6 ± 8 |
| Ugni Blanc | 26 ± 12 [4, 49] | 41 | 20 ± 28 |
| Melon | 21 ± 10 [2, 49] | 39 | 6 ± 4 |
| Chenin | 18 ± 8 [1, 40] | 38 | 5 ± 5 |
| Gewurztraminer | 28 ± 9 [2, 49] | 28 | 9 ± 10 |
| Pinot Auxerrois | 32 ± 8 [11, 49] | 24 | 8 ± 8 |
| Riesling | 28 ± 9 [10, 49] | 24 | 8 ± 10 |

The model consisted of two hierarchical structured components: one describing the prevalence of esca at plot scale (presence of esca, denoted 1, or absence, denoted 0), and the other the incidence of esca conditional on its occurrence (i.e. given that the plot prevalence is 1). A Bernoulli distribution with a logit link was used for the prevalence component. For the incidence component, the response variable was the number of vines with esca leaf symptoms divided by the total number of vines observed. For this component, we aimed to establish a binomial model conditional on there being at least one vine with esca symptoms in the plot. Thus, a zero-inflated binomial distribution parameterised to exclude zero with a logit link was used for the incidence. Both the prevalence and incidence components included cultivar and plot identity as independent random effects and the year as an autoregressive process of order 1 (AR1). These effects account for interannual dependence arising from underlying meteorological or environmental factors and agricultural practices affecting esca symptoms. Finally, the incidence component included plot age as an AR1 process specific to each cultivar. We describe below the “full” model including all explanatory variables. Intermediate models with fewer explanatory variables were also estimated and compared to the full model on the basis of two information criteria: the deviance information criterion and the widely applicable information criterion (DIC and WAIC) (Table 2).

**Table 2:** Deviance information criterion (DIC) and widely applicable information criterion (WAIC) for the eight model structures considered. Number of vines monitored per plot: n_vines_.

| Model ID | Prevalence model | Incidence model | DIC | WAIC |
| --- | --- | --- | --- | --- |
| M8 “Full” | plot + cultivar + year + $n_{\text{vines}}$ | plot + cultivar + year + age x cultivar | 37842 | 64418 |
| M7 | plot + cultivar + year + $n_{\text{vines}}$ | plot + cultivar + year + age | 40272 | 80397 |
| M6 | plot + cultivar + year + $n_{\text{vines}}$ | plot + cultivar + year | 39771 | 111086 |
| M5 | plot + cultivar + year + $n_{\text{vines}}$ | plot + cultivar | 44832 | 142323 |
| M4 | plot + cultivar + year + $n_{\text{vines}}$ | plot | 44871 | 142351 |
| M3 | plot + cultivar + year + $n_{\text{vines}}$ | 1 | 132737 | 121358 |
| M2 | plot + cultivar + year | 1 | 132736 | 121357 |
| M1 | plot + cultivar | 1 | 132813 | 121436 |
| M0 | plot | 1 | 132913 | 121497 |

More specifically, the variables in the model are as follows: let *y_i_* denote the esca prevalence of observation *i* (*i* = 1, …, 5161), where the disease is either observed (value *y_i_* = 1) or not observed (value *y_i_* = 0). Let *n_i_* denote the number of plants with esca symptoms among the *n_i_^tot^* monitored for this same observation *i*. We will consider the following explanatory variables associated with observation *i*. First, the variable *plot(i)* denotes the identity of the plot (554 levels for prevalence and 546 levels for incidence) on which observation *i* was performed; *cultivar(i)* denotes the cultivar (12 levels, see Table 1) on the plot corresponding to observation *i*. The variable *year(i)* for prevalence denotes the year of monitoring (indexed by *t* = 2003, …, 2022). Finally, *age*(*i*) denotes plot age, a variable considered only for the incidence component (indexed by *z* = 1, …, 49) being specific to each cultivar.

The first component (prevalence) of the model can be written as follows:

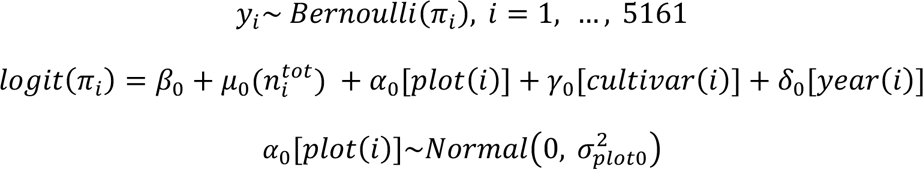

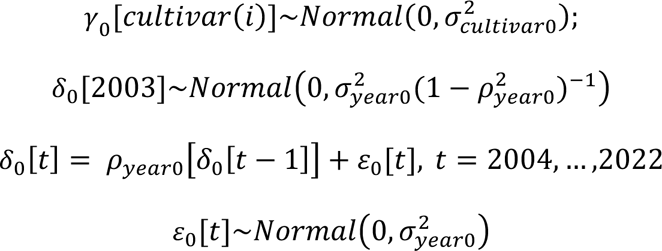

ZIB^+^ is a modified version of the zero-inflated binomial distribution supported from 1 to n_i_^tot^ (exclusion of the 0 value), *S* being the subset of index *i* associated when esca prevalence = 1 (4476 observations).

The second component (incidence) of the model can therefore be written as follows:

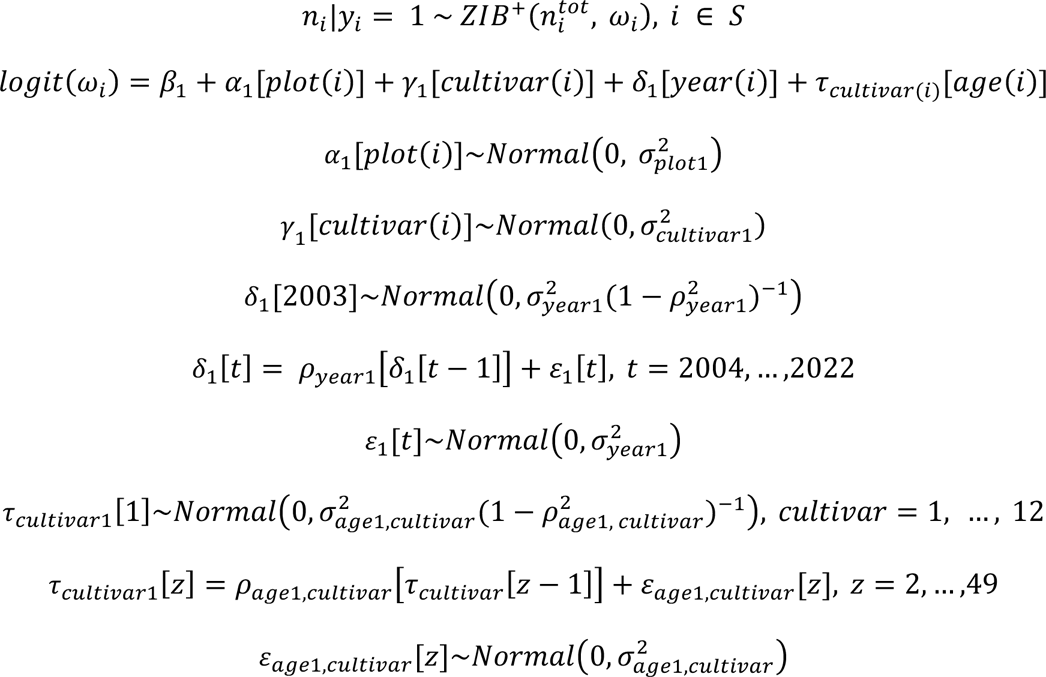

In the model, β_0_ and β_1_ are the intercepts of prevalence and incidence, respectively. The parameter *µ_0_* takes into account the different numbers of plants monitored per plot (note that *n_i_^tot^* was standardised to ensure robust estimation). The autoregressive parameters *ρ_year0_, ρ_year1_, ρ_age1,cultivar_* (all of which have absolute values < 1) correspond to the correlation coefficients for the prevalence (*ρ_year0_* for year) or incidence (*ρ_year1_* for year and *ρ_age1,cultivar_* for plot age for a given cultivar) of esca between two consecutive time points. They indicate the extent to which the temporal trend in a given time step *t* depends on the trend from the previous time step *t*-1. Typically, estimates close to 0 indicate the absence of a significant temporal trend. By contrast, the closer the estimates are to 1, the more similar the temporal trends between two consecutive time steps (Zuur et al., 2017). The parameters *σ^2^_year0_, σ^2^_year1_, σ^2^_age1,cultivar_* define the precision of the AR1 process, with lower values corresponding to smoother temporal trends. The model was fitted in a Bayesian framework by the integrated nested Laplace approximation (INLA) method (Rue et al., 2009, 2017) with R (version 4.2.1). Bayesian inference requires the specification of prior distributions for the model parameters and hyperparameters, and we used the default internal vague priors recommended in R-INLA. Specifically, the different *σ^2^_plot0_, σ^2^_cultivar0_, σ^2^_plot1_, σ^2^_cultivar1_* values for the independent random effects and the *σ^2^_year0_, σ^2^_year1_, σ^2^_age1,cultivar_* values of the AR1 process are assigned a log-gamma distribution with parameters 1 and 0.00005. For the cultivar and year effects, we then calculated the probability of direction (*pd*), which varies between 0.5 and 1 and indicates the probability that a parameter, as defined by its posterior distribution, is either strictly positive or negative. This *pd* is the proportion of the posterior distribution that has the same sign as the median of the posterior distribution (Makowski et al., 2019) and can be used to identify trends in the distribution of the parameter. Finally, we compared fitted and observed values to evaluate model fit. For the probability of presence (prevalence), we used ROC curve analyses (Hoo et al., 2017, PRROC R package, Grau et al., 2015). For disease incidence, we calculated Pearson’s R coefficient for the correlation between the mean of the fitted values and observed incidence (function *cor.test* package *stats*).

## III) Results

### 1) Prevalence and incidence of esca and Eutypa dieback leaf symptoms in France between 2003 and 2022, by cultivar and wine region

#### 1.1) Variability of the prevalence and incidence of esca leaf symptoms

Only 10% of the 2,075 plots monitored over the 2003-2022 period contained no symptomatic vines (1,861 of 2,075 plots contained at least one vine presenting symptoms). The annual prevalence of esca ranged from 54% in 2010 to 84% in 2017, with a mean of 74 ± 2% (mean ± SEM, standard error of the mean) plots containing at least one symptomatic plant (Figure 3a). During the monitoring period, for all years considered together, the prevalence of esca in the regions ranged from 100% of the plots containing at least one vine with esca (Corse and Jura & Savoie) to 78% (Champagne) (data not shown).

Including plots without esca (prevalence = 0), the mean annual incidence of esca over the 2003-2022 period was 3.1 ± 0.1% (mean ± SEM). The lowest mean annual incidence of esca was observed in 2003 whereas the highest incidence was observed in 2012, 2014, and 2017 (Figure 3a). The annual incidence of esca increased between 2003 and 2008 and then fluctuated until 2022. If we excluded plots with no vines presenting esca symptoms in a given year (prevalence = 0), the mean annual incidence increased to 4.2 ± 0.1% (mean ± SEM, Figure S2).

We observed considerable variability in the incidence of esca leaf symptoms between cultivars. The mean incidence by cultivar ranged from 0.6% to 10.6% (Figure 5a). The Trousseau cultivar had the highest incidence, followed by Savagnin and Ugni Blanc. Incidence was lowest for Meunier, followed by Pinot Noir and Syrah.

**Figure 5:**
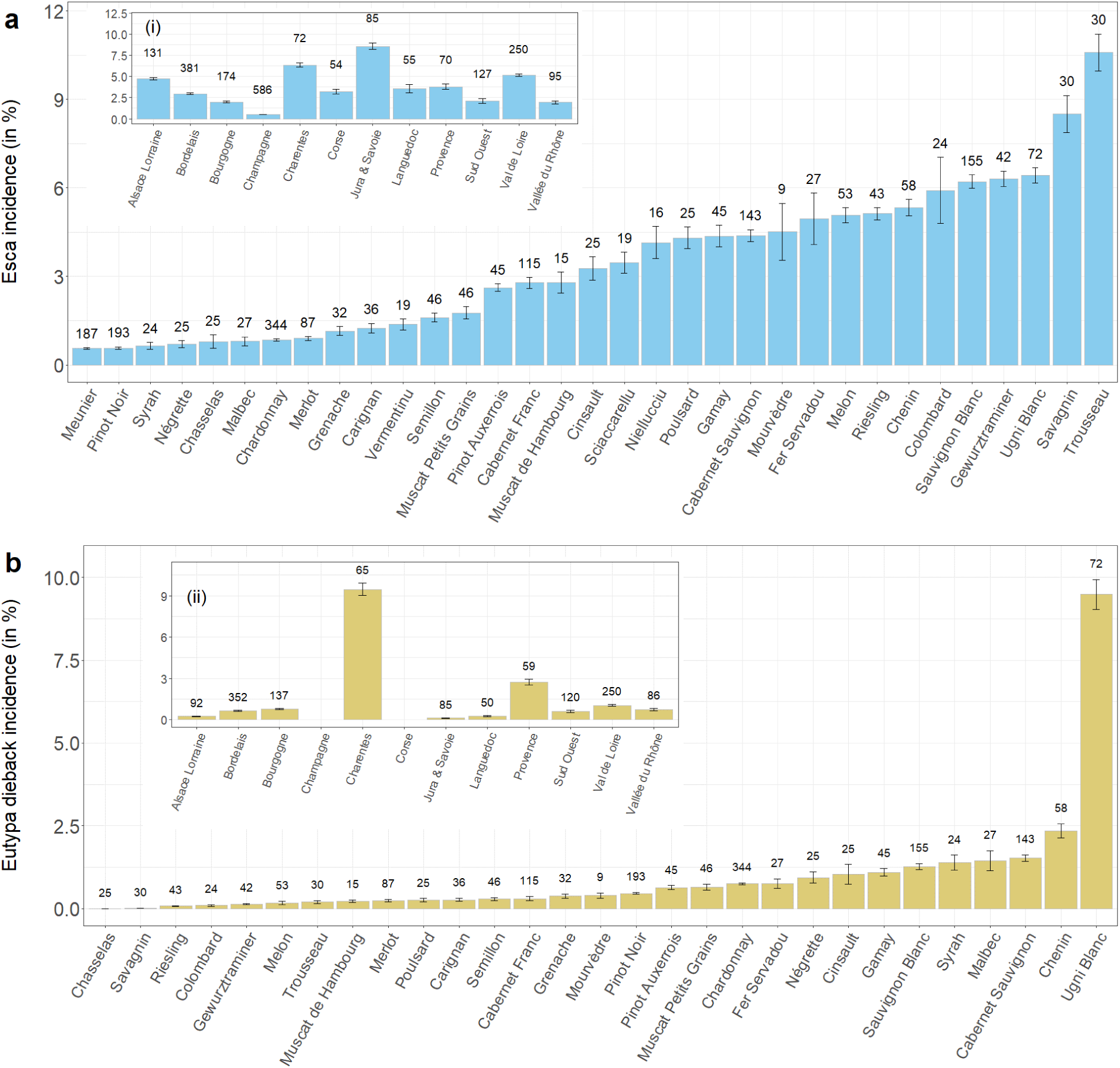
The mean incidence of esca and Eutypa dieback leaf symptoms by cultivar and wine region (inset, i : esca; ii : Eutypa dieback) in France over the 2003-2022 period. Numbers represent the number of plots monitored for the incidence of (a) esca (blue bars) and (b) Eutypa dieback (mustard bars). The error bars represent the standard error of the mean.

The incidence of esca also varied between wine regions, the highest incidence being recorded in Jura & Savoie and the lowest in Champagne(Figure 5a(i)). The incidence of esca by year, region and cultivar was not correlated with the number of plots monitored (Pearson’s r, *p* > 0.13)

#### 1.2) Variability of the prevalence and incidence of Eutypa dieback

In total, 1,296 plots were monitored for Eutypa dieback over the 2003-2022 period, and 31% were Eutypa-free (895 of the 1,296 plots contained vines displaying Eutypa dieback). The annual percentage of plots with Eutypa dieback (i.e, the prevalence) ranged from 20% in 2016 to 61% in 2005, with a mean value of 41 ± 3% (mean ± SEM) (Figure 3b). During the monitoring period, the regional prevalence of Eutypa dieback ranged from 96% in Charentes to 40% in the Sud Ouest region.

If plots without Eutypa dieback (prevalence = 0) were included in the calculation, then the mean observed incidence of Eutypa dieback was 1.4 ± 0.1% (mean ± SEM) (Figure 3b). The incidence of Eutypa dieback tended to decrease from 2003 to 2022 (Figure 3b). Specifically, we observed a decrease from 2003 to 2008 and then a steady decrease since 2013 (except for 2020). The incidence of Eutypa dieback was highest in 2003. In addition, 2017 and 2021 were the years in which the incidence of Eutypa dieback was lowest. Finally, if plots in which none of the vines were affected by Eutypa dieback (prevalence = 0) in a given year were excluded, the mean incidence increased to 3.3 ± 0.1% (mean ± SEM, Figure S2).

The variability of Eutypa dieback incidence between cultivars was high, with values ranging from 0% to 9.5% (Figure 5b). The region with the highest Eutypa dieback incidence was Charentes and with the lowest, Jura & Savoie and Alsace Lorraine (Figure 5b(ii)). The incidence of Eutypa dieback by year, region and cultivar was not correlated with the number of plots monitored (Pearson’s r, *p* > 0.30). Finally, there was a weak but significant correlation between the incidences of esca and Eutypa dieback assessed by plot/year observation (Pearson’s r = 0.08, *p* < 0.001).

### 2) Deciphering the effects of cultivar, year, and plot age on the prevalence and incidence of esca leaf symptoms

The two model selection criteria considered (DIC and WAIC) supported the full model (Table 2), suggesting that all the explanatory variables considered had a significant impact on esca dynamics. We therefore consider only the full model hereafter. The model fit was satisfactory for both the prevalence and incidence components. Specifically, the area under the ROC curve (AUC) for esca prevalence was 0.93. Pearson’s r for the correlation between adjusted and observed values of esca incidence was 0.83 (*p* < 0.001).

The model included the number of vines monitored as a covariable to account for the sampling effort in each plot used to determine incidence. As expected, the mean effect was positive (0.33), confirming that increasing the number of plants inspected in a plot increases the likelihood of at least one plant being symptomatic. The associated probability of direction was 0.95 (µ_0_, Table 3). The identity of the plot had a significant impact on both the prevalence and incidence of esca symptoms. For prevalence, 13% of the individual plot effects were negative (i.e. associated with a q-97.5% < 0), < 1% were positive (q-2.5% > 0) and the remaining 87% were associated with individual effects including zero (the mean value) in their 95% credible intervals. For incidence, 25% of the plots were negative, 30% were positive and the remaining 45% were associated with individual effects including zero in the 95% credible interval. These values correspond to the percentage of plots characterised by an esca prevalence or incidence lower than, higher than or not significantly different from the overall mean, respectively.

**Table 3:** Summary statistics for the parameters of the full model of the prevalence and incidence of esca leaf symptoms. For both model components (prevalence and incidence for plots with a prevalence of 1), the table summarises the posterior mean, standard deviation (SD), 0.025, 0.5, 0.975 quantiles (quant.) and mode for intercepts (β_0_: intercept for esca prevalence; β_1_: intercept for esca incidence), the number of vines monitored (µ_0_), the variance of the independent random effects associated with cultivars and plots (*σ^2^_cultivar0_, σ^2^_cultivar1_, σ^2^_plot0_ and σ^2^_plot1_*), the correlation coefficients and variance of the AR1 process associated with year (*ρ_year0_* and *ρ_year1_*) and the correlation coefficients and variance of the AR1 process associated with age for each cultivar (*ρ_age1,cultivar_* and *σ^2^_age1,cultivar_* to be detailed).

|  | Mean | SD | 0.025quant. | 0.5quant. | 0.975quant. | Mode |
| --- | --- | --- | --- | --- | --- | --- |
| Esca prevalence |  |  |  |  |  |  |
| $\beta_0$ | 2.75 | 0.47 | 1.83 | 2.75 | 3.69 | 2.74 |
| $\mu_0$ | 0.33 | 0.20 | -0.05 | 0.33 | 0.73 | 0.33 |
| $\sigma^2_{cultivar0}$ | 0.90 | 0.40 | 0.35 | 0.83 | 1.88 | 0.70 |
| $\sigma^2_{plot0}$ | 0.36 | 0.04 | 0.29 | 0.36 | 0.44 | 0.36 |
| $\rho_{year0}$ | 0.64 | 0.16 | 0.25 | 0.67 | 0.89 | 0.72 |
| $\sigma^2_{year0}$ | 5.11 | 2.40 | 1.84 | 4.65 | 11.08 | 3.82 |
| Esca incidence (when prevalence > 0) |  |  |  |  |  |  |
| $\beta_1$ | -3.91 | 0.20 | -4.41 | -3.90 | -3.46 | -3.88 |
| $\sigma^2_{cultivar1}$ | 3.53 | 2.42 | 0.94 | 2.89 | 9.95 | 2.01 |
| $\sigma^2_{plot1}$ | 1.25 | 0.09 | 1.09 | 1.25 | 1.43 | 1.25 |
| $\rho_{year1}$ | 0.30 | 0.24 | -0.20 | 0.32 | 0.71 | 0.36 |
| $\sigma^2_{year1}$ | 12.58 | 4.35 | 5.87 | 11.98 | 22.79 | 10.84 |
| $\rho_{age1,Cabernet Franc}$ | 0.80 | 0.09 | 0.59 | 0.82 | 0.93 | 0.85 |
| $\sigma^2_{age1,Cabernet Franc}$ | 2.81 | 1.32 | 1.03 | 2.54 | 6.12 | 2.09 |
| $\rho_{age1,Cabernet Sauvignon}$ | 0.95 | 0.03 | 0.87 | 0.95 | 0.98 | 0.96 |
| $\sigma^2_{age1,Cabernet Sauvignon}$ | 0.72 | 0.38 | 0.24 | 0.64 | 1.68 | 0.50 |
| $\rho_{age1,Chardonnay}$ | 0.80 | 0.10 | 0.55 | 0.82 | 0.93 | 0.85 |
| $\sigma^2_{age1,Chardonnay}$ | 2.62 | 1.36 | 0.89 | 2.32 | 6.09 | 1.83 |
| $\rho_{\text{age1,Chenin}}$ | 0.82 | 0.09 | 0.60 | 0.83 | 0.94 | 0.87 |
| $\sigma^2_{\text{age1,Chenin}}$ | 2.59 | 1.27 | 0.87 | 2.35 | 5.77 | 1.89 |
| $\rho_{\text{age1,Gewurztraminer}}$ | 0.43 | 0.16 | 0.09 | 0.44 | 0.71 | 0.46 |
| $\sigma^2_{\text{age1,Gewurztraminer}}$ | 1.33 | 0.39 | 0.72 | 1.28 | 2.23 | 1.19 |
| $\rho_{\text{age1,Melon}}$ | 0.77 | 0.13 | 0.45 | 0.80 | 0.94 | 0.85 |
| $\sigma^2_{\text{age1,Melon}}$ | 6.04 | 2.92 | 2.14 | 5.46 | 13.39 | 4.42 |
| $\rho_{\text{age1,Merlot}}$ | 0.74 | 0.13 | 0.44 | 0.76 | 0.93 | 0.81 |
| $\sigma^2_{\text{age1,Merlot}}$ | 1.62 | 0.80 | 0.52 | 1.47 | 3.59 | 1.18 |
| $\rho_{\text{age1,Pinot Auxerrois}}$ | 0.66 | 0.14 | 0.33 | 0.69 | 0.88 | 0.73 |
| $\sigma^2_{\text{age1,Pinot Auxerrois}}$ | 4.83 | 2.20 | 1.82 | 4.41 | 10.30 | 3.67 |
| $\rho_{\text{age1,Pinot Noir}}$ | 0.91 | 0.05 | 0.80 | 0.92 | 0.97 | 0.94 |
| $\sigma^2_{\text{age1,Pinot Noir}}$ | 0.97 | 0.50 | 0.30 | 0.87 | 2.22 | 0.68 |
| $\rho_{\text{age1,Riesling}}$ | 0.80 | 0.10 | 0.55 | 0.82 | 0.94 | 0.86 |
| $\sigma^2_{\text{age1,Riesling}}$ | 6.85 | 3.45 | 2.27 | 6.15 | 15.53 | 4.90 |
| $\rho_{\text{age1,Sauvignon Blanc}}$ | 0.82 | 0.08 | 0.64 | 0.84 | 0.94 | 0.86 |
| $\sigma^2_{\text{age1,Sauvignon Blanc}}$ | 2.17 | 0.99 | 0.74 | 2.01 | 4.54 | 1.67 |
| $\rho_{\text{age1,Ugni Blanc}}$ | 0.90 | 0.06 | 0.76 | 0.91 | 0.97 | 0.93 |
| $\sigma^2_{\text{age1,Ugni Blanc}}$ | 5.37 | 2.89 | 1.61 | 4.77 | 12.66 | 3.67 |

Cultivar had a significant impact on the prevalence of esca symptoms but a much weaker effect on esca incidence. Indeed, cultivar was significant if WAIC was used (64436 without the cultivar effect vs. 64418 for the full model) but not with the DIC (37835 without the cultivar effect vs. 37842 for the full model, Table 2). The individual effects of each of the 12 cultivars considered are shown in Figure 6. Chardonnay, Merlot and Pinot Noir all had a prevalence of esca below the prevalence above the overall mean (q-2.5% > 0) (Figure 6a). No such differences were observed for the incidence of esca. The 12 estimated 95% credible intervals included zero, meaning that esca incidence did not differ between cultivars, provided that esca was observed (prevalence > 0; Figure 6b). However, the probability of direction (*pd*) showed that cultivars such as Gewurztraminer, Riesling and Melon had a strong tendency to have a high incidence whereas Merlot and Chardonnay had a low incidence (*pd* > 0.90, Figure 6a, b).

**Figure 6:**
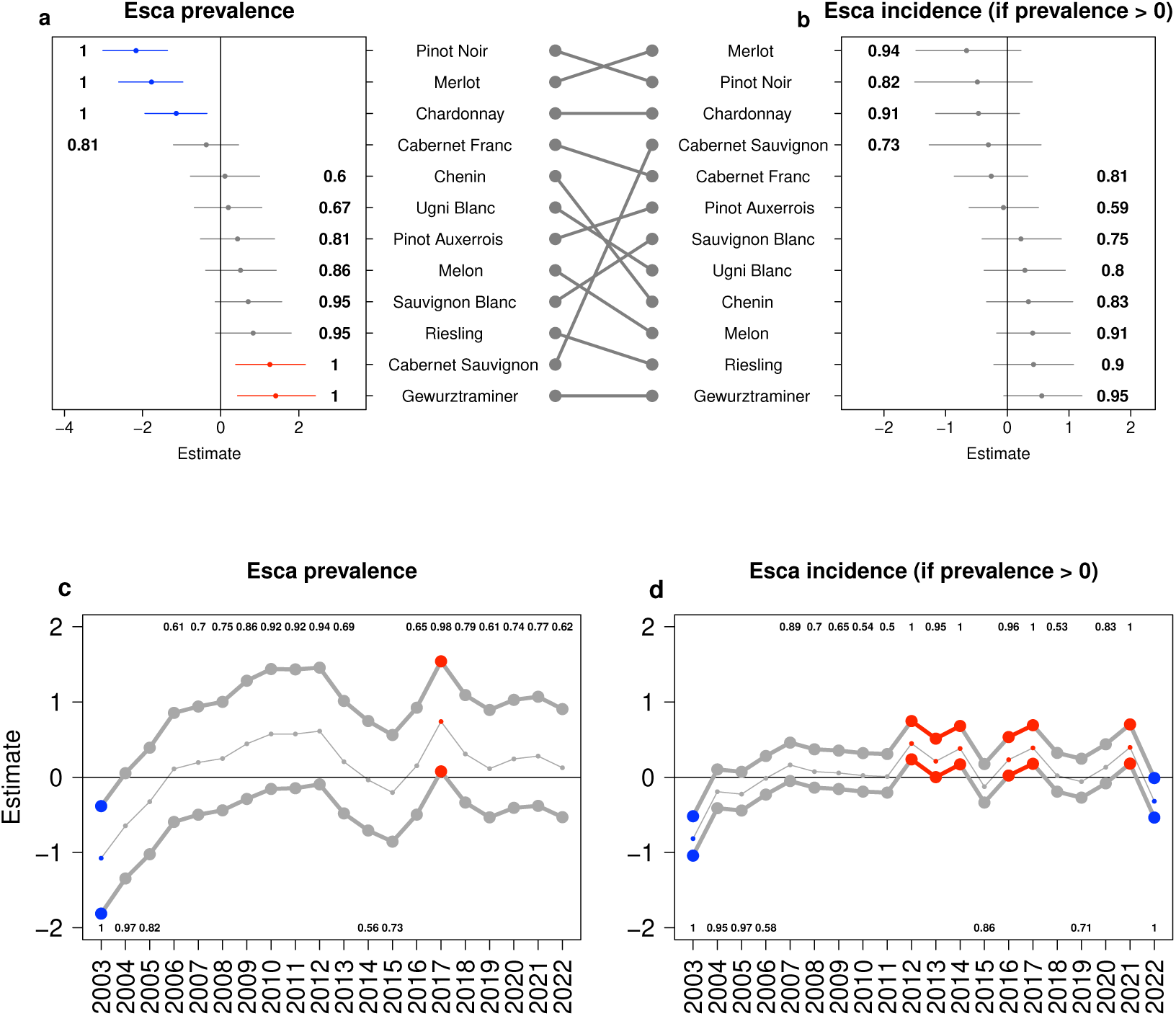
Effects of cultivar (upper panels, means in descending order) and year (lower panels) on esca prevalence (left panels) or esca incidence if esca was observed in the plot (right panels). In the upper panels, black points represent the posterior median (50th percentiles, middle) and horizontal lines the lower (2.5th percentiles) and upper (97.5th percentiles) limits of the credible interval for the independent random effects associated with each cultivar. The segments between the upper panels (a and b) show the relationship between the estimated prevalence and incidence of the different cultivars. The lower panels display the year effects for esca prevalence and incidence, as fitted with an autoregressive model of order 1. Estimates in blue correspond to years associated with significantly negative effects (97.5th percentile < 0) relative to the mean (0), whereas those in red correspond to years with significantly positive effects (2.5th percentile > 0). The values shown correspond to the probability of direction for each cultivar (see the materials

Unlike the effect of cultivar, the effect of year did not affect the prevalence of esca symptoms, but did affect their incidence. However, the 95% credible interval [0.25, 0.89] of the hyperparameters *ρ_year0_* of the AR1 process modelling the effect of year on esca prevalence suggests that esca prevalence is strongly correlated across successive years. No such significant correlation was detected for esca incidence (the 95% CI of *ρ_year1_* was [-0.20, 0.71]) (Table 3; Figure 6c, d). An analysis by year showed that the prevalence or incidence (if esca was recorded in the plot) of esca was significantly lower than the mean value (=0) in the first year (2003) and that esca incidence was lower in 2022. The prevalence and incidence of esca tended to be below the mean value in 2004, as was the incidence in 2005 (probability of direction, *pd* > 0.95). In 2017, the prevalence and incidence of esca were significantly higher than the mean. Moreover, esca incidence has remained above the mean value since 2012, including the years 2013, 2014, 2016, 2017, and 2021. However, *pd* values showed that between 2010 and 2012, prevalence tended to be high (pd > 0.92, as shown in Figure 6c, d).

The hyperparameters *ρ_age1,cultivar_* of the AR1 process, modelling the effect of plot age on esca incidence for each cultivar, were strictly positive, suggesting a positive correlation between plot age and incidence (Table 3). A pattern of the effect of age on esca incidence common to all cultivars is emerging. Indeed, in most cultivars, esca symptoms peak at an intermediate age. With the exception of Merlot, all cultivars had an incidence of esca above the mean value for plots aged between 10 and 40 years (in red, Figure 7). Conversely, the youngest plots (under 10 years old) sown with these cultivars had an esca incidence below the mean value (in blue, Figure 7). Finally, Cabernet Franc, Cabernet Sauvignon, Gewurztraminer, Merlot and Pinot Noir had incidences significantly below the mean value after the age of 30 years. Overall, when age had a significant positive effect on esca incidence (when esca prevalence > 0), esca occurred at ages between nine and 45 years, with a peak of susceptibility between nine and 24 years, depending on the cultivar (Figure 8). Even though the cultivars with the lowest incidence had a narrower age range for peak incidence (Cabernet Franc, Cabernet Sauvignon, Chardonnay, Pinot Noir), a pattern remained in the relationship between cultivar susceptibility (from left to right, according to modelling, Figures 6 and 8) and the age ranges for susceptibility to esca and peak incidence.

**Figure 7:**
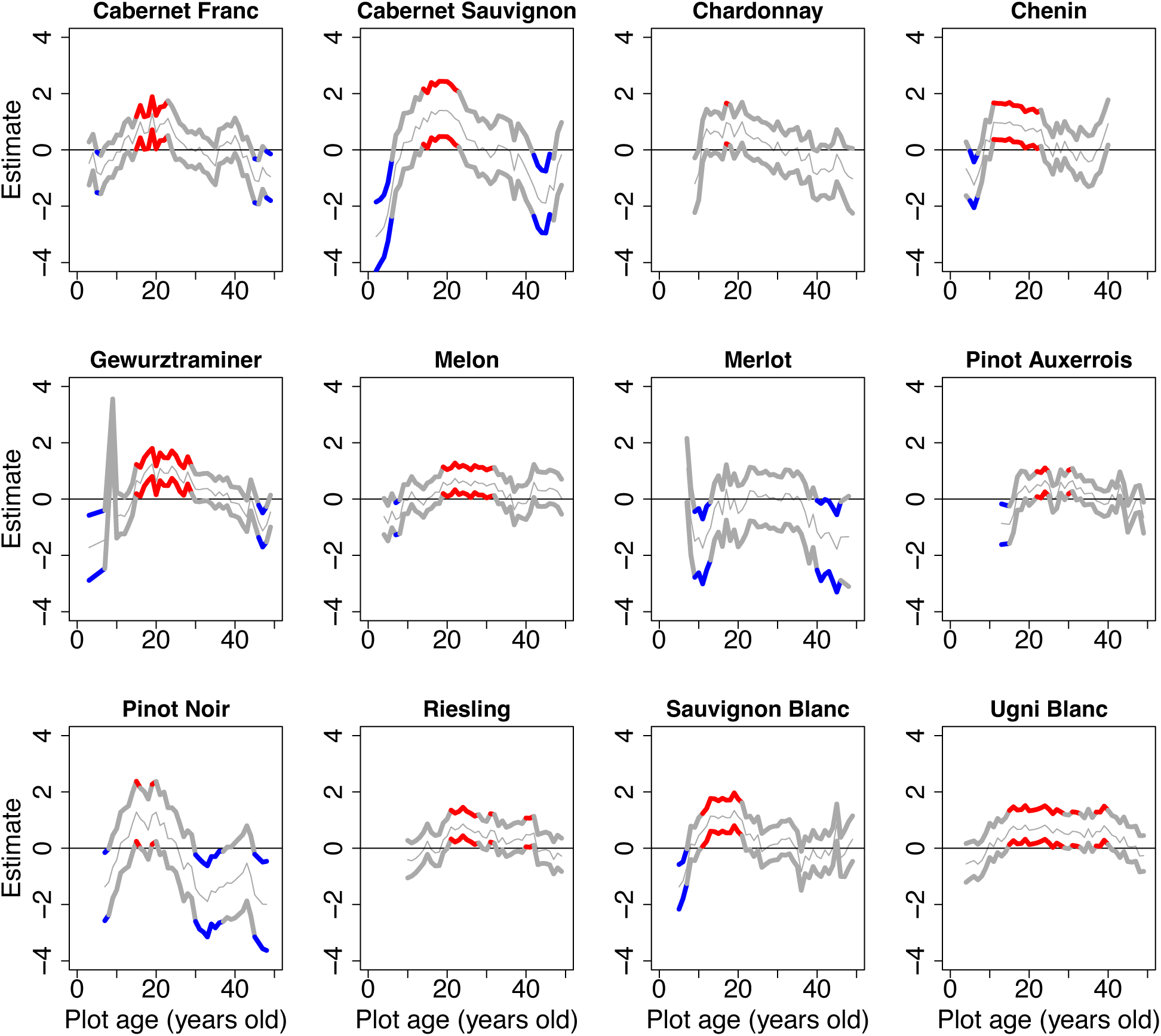
Impact of plot age (in years), by cultivar, on standardised estimates of the esca incidence for the plots in which esca was observed. The posterior median (50th percentiles, middle), lower (2.5th percentiles) and upper (97.5th percentiles) limits of the credible interval are shown. Estimates in blue correspond to ages associated with significantly negative effects (97.5th percentile < 0) relative to the mean (0) whereas those in red correspond to ages associated with significantly positive effects (2.5th percentile > 0). Estimates are shown in colour if the findings for at least two consecutive years are significant.

**Figure 8:**
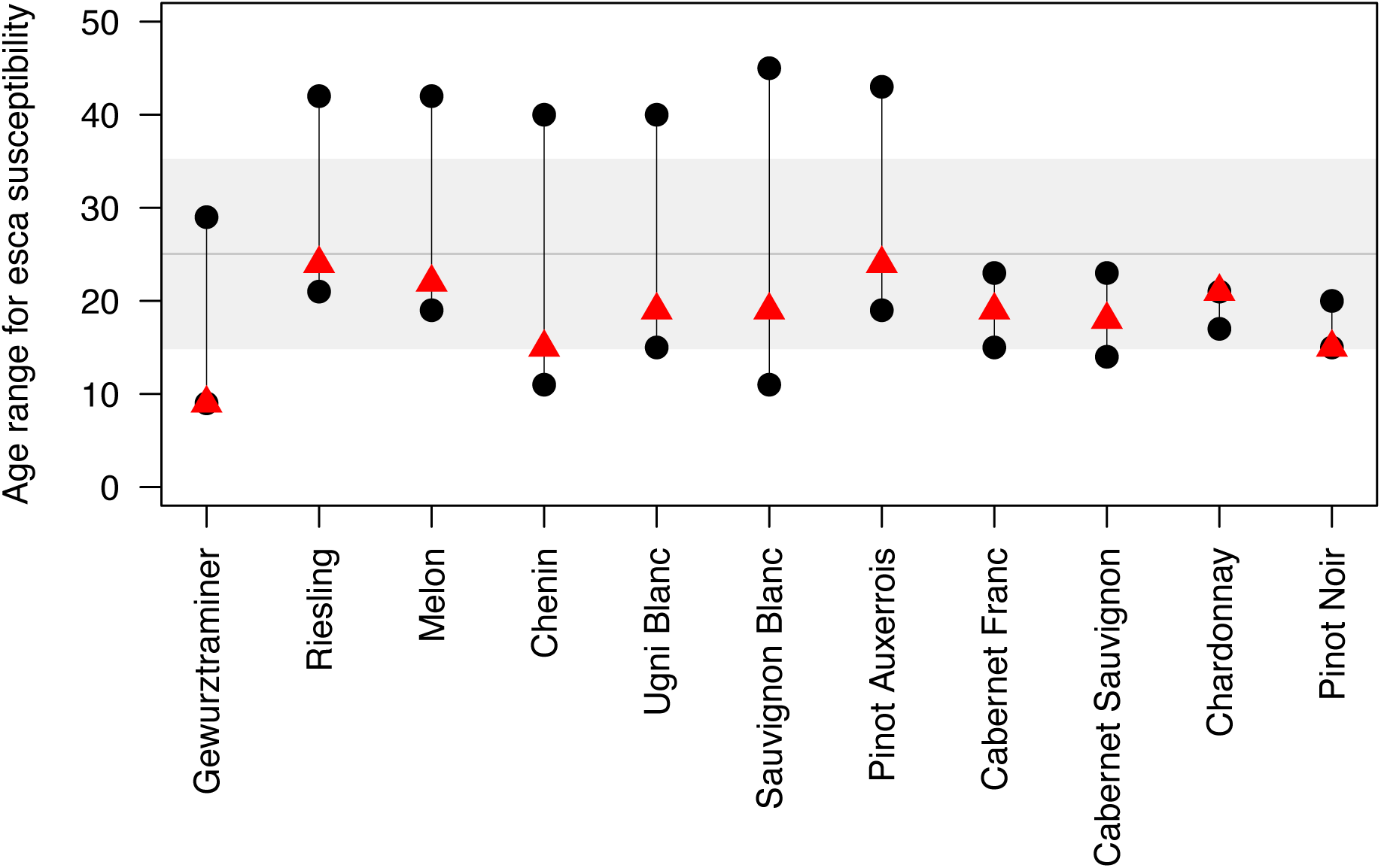
Range of ages during which vines are susceptible to esca (black points) and peak (red triangle) susceptibility for cultivars with significant and positive results between the first and the last plot age. The cultivars are ordered by estimated mean incidence and only cultivars with an age significantly and positively above the mean of 0 are shown (Figure 6b, Merlot is therefore not represented). The wide horizontal grey band represents the standard deviation from the mean, which is represented by the horizontal dark grey bar, for all cultivars combined.

## IV) Discussion

We compiled and homogenised regional and historical databases to obtain a national quantification of the prevalence and incidence of leaf symptoms of esca and Eutypa dieback in France between 2003 and 2022. Despite considerable variability between years and cultivars, the incidence and prevalence of esca leaf symptoms appear to be stable when looking at the raw data, whereas the incidence of Eutypa dieback tended to decrease over the 2003-2022 period. We used a hierarchical Bayesian model to decipher the responses of two components — the prevalence (percentage of symptomatic plots, i.e. plots with at least one symptomatic plant) and the incidence (percentage of symptomatic plants per plot, when prevalence > 0) of esca disease — to the effects of year, cultivar, and plot age. This framework is compatible with the objectives of surveillance programmes in applied plant pathology, which often aim to estimate disease occurrence at national, regional, and local scales. The framework developed here takes into account disease occurrence at two nested spatial scales (a set of plots within each agricultural region and a set of plants within each plot). In our case study, focusing on the prevalence of esca leaf symptoms is particularly relevant when comparing a wide range of cultivars with different levels of susceptibility, including some for which no leaf symptoms were reported on a large number of plots. Once the presence of leaf symptoms is confirmed in a plot, analyses of the variation of incidence are then required to identify the factors driving leaf symptom expression, as esca is characterised by fluctuating expression between years and plots. This approach showed that since the incidence of esca leaf symptoms was significantly higher than the mean value in six of the years since 2003, all occurring after 2012.

This study highlights the strong annual variability of esca incidence. Analysis of the whole database showed an increase in the annual mean incidence of esca from 2003 (1.6%) to 2008 (3.9%). This trend was already reported by Bruez et al., (2013). However, our longer-term monitoring revealed that this trend ceased after 2008. More specifically, the upward trend seems to have continued after 2008 for certain cultivars, such as Ugni Blanc and Riesling in particular, and their associated regions (Figure S3 and S4), whereas the opposite was observed for Chardonnay and Meunier (Figure S4). Statistically, when esca is observed in a plot, leaf symptoms may be expressed over a number of years. The differences between years may reflect climatic variations. Indeed, the lowest esca incidence was recorded in 2003, a year characterised by a very intense drought and heatwave (Chuine et al., 2004). Other years of low esca incidences seem to coincide with hot summers (e.g., 2015, 2019, 2022, https://climate.copernicus.eu/europe-continued-swelter-july, site visited on February 22^nd^, 2024). In the literature, it has been suggested that higher maximum temperatures between June and July (leading to drier environments) are associated with a lower incidence of esca leaf symptoms and similarly the incidence of esca leaf symptoms seems to be positively correlated with precipitation levels, as high rainfall levels during July tend to intensify leaf symptom development (Calzarano et al., 2018; Serra et al., 2018). Indeed, water availability has been shown to affect esca leaf symptom development as, under controlled conditions, intense drought inhibits esca leaf symptom onset, suggesting an antagonistic interaction between drought and esca pathogenesis and a key role for plant water status and climatic conditions (Bortolami et al., 2021). Vascular diseases with similar characteristics to esca are rare. One example is provided by a trunk disease of kiwifruits in which there is a significant relationship between temperature and leaf symptoms (Di Marco and Osti, 2008).

An analysis of the whole database suggested that, unlike esca, Eutypa dieback decreased steadily in incidence during the 2003-2022 period. These patterns were observed across the most observed cultivars, years, and wine regions (Figure S5 and S6). The difference between the development of esca and Eutypa dieback symptoms over time may partly reflect the nature of these diseases. Esca is generally caused by a complex community of fungi, whereas foliar symptoms of Eutypa dieback is likely caused by a single fungus, *Eutypa lata*. Climate can affect the expression of Eutypa dieback (Sosnowski et al., 2007), but its gradual decrease in frequency over the years suggests that climate has a relatively limited impact. The observed gradual decrease in Eutypa dieback frequency may be due to the use of effective control methods, such as appropriate pruning, wound protection and the removal of infected wood from vines (Lecomte et al., 2006), or to a decrease in pathogen aggressiveness (Molyneux et al., 2002). In any case, Eutypa dieback has been relegated to the status of a secondary disease, attracting less attention from field technicians, as indicated by the decrease in the number of plots monitored for this disease over the years (Figure S5 and S6; except for Ugni Blanc).

Our findings revealed considerably variability in the incidence of esca between the 36 cultivars monitored in the whole database. Cultivar differences in esca incidence have already been documented in France (Bruez et al., 2013; Gastou et al., 2024), Italy (Murolo and Romanazzi, 2014) and Spain (Chacón-Vozmediano et al., 2021). In these studies, Cabernet Sauvignon and Sauvignon Blanc were generally identified as much more susceptible than Merlot and Chardonnay (Murolo and Romanazzi, 2014; Chacón-Vozmediano et al., 2021; Gastou et al., 2024). However, our statistical analysis revealed a lack of significant difference between the cultivars if esca incidence was calculated only for plots with at least one symptomatic plant. Interestingly, the prevalence of esca was significantly higher than the mean for two cultivars and lower than the mean for three cultivars. Moreover, the cultivars with the highest prevalence of esca also had the highest estimated incidence values (Figure 6). Only Cabernet Sauvignon displayed contrasting levels of esca prevalence and incidence, as most of the plots had at least one symptomatic plant but an intermediate incidence. Several confounding factors could potentially cause the cultivar effect highlighted in our analysis. Typically, some viticulture practices (e.g. the use of certain rootstocks or types of pruning) may be specific to a particular cultivar and wine region. Such factors are known to affect susceptibility to esca (Lecomte et al., 2018). It was not possible to investigate these confounding factors here, but the observed differences in esca incidence between the cultivars studied here were correlated with the range of susceptibility of the same cultivars quantified over several years in a common garden vineyard (Gastou et al., 2024).

Our results revealed a major impact of plot age on esca incidence, which was lower in younger vineyards (less than 10 years old) than in older ones. This age effect has also been observed in several field monitoring programmes (Fulchin et al., 2019; Fussler et al., 2008; Pollastro et al., 2000). The underlying mechanisms remain to be explored. We could hypothesize that certain traits known to differ with plant age, such as plant metabolism, wood properties, and pathogen or endophyte communities, are involved. As suggested by Fischer and Peighami-Ashnaei (2019), older plants might be more susceptible to disease due to cumulated wounds over time, as cultivated vines are pruned annually. Older plants may therefore have undergone more infection cycles than younger plants. In addition, older plants tend to accumulate more endophytes in their tissues than younger plants (Dissanayake et al., 2018). They also have different fungal communities (Bruez et al., 2016) and may undergo microbiome modifications during the course of their lifetime (Bettenfeld et al., 2020; Fournier et al., 2022). Indeed, many endophytic fungi are pathogens and can induce plant dieback (e.g., Úrbez-Torres et al., 2009).

Interestingly, the effect of age on the incidence of esca differed between cultivars. There was a trend towards a relationship between cultivar susceptibility (i.e. esca incidence) and the age range during which the cultivar was most susceptible (i.e. highest incidence), the least susceptible cultivars had a narrower age range for susceptibility. In four of the 12 cultivars studied, esca incidence decreased significantly once a certain age was reached, this age depending on the cultivar but generally being greater than 30 years. This pattern may reflect a higher rate of plant mortality in older plots, with the most tolerant plants remaining, the others generally being replaced by new replanted vines. Indeed, in our database, we observed a slight gradual ageing of the plots from 2003 to 2014 and an increase in the diversity of plot ages (monitoring of new younger plots to replace the older ones) starting in 2015 (Figure 3). If we wish to study the effect of vine age on the incidence of esca more precisely, we need to be able to collect the age of the vines rather than the age of the plot (based on the year of first plantation). Alternatively, ontogenic resistance (Ficke et al., 2002) could be involved, leading to a lower susceptibility to the development of esca leaf symptoms in the oldest vines, but little is known about this process in wood diseases. Secondary metabolism may change qualitatively or quantitatively with plant ageing (Haffner et al., 1991), potentially accounting for the persistence of long-lived perennial plants (Cui et al., 2024). With regard to other diseases, a higher incidence of Botryosphaeriaceae (Carlucci et al., 2013; Gubler et al., 2005) and *Phaeoacremonium sp.* (Carlucci et al., 2013, 2015) has been observed in older grapevines and olive trees. Similarly, an increase in branch canker and dieback incidence with the age of the tree has been observed in avocado orchards (Valencia et al., 2022). By contrast, for kiwifruit (Di Marco and Osti, 2008), no correlation was found between plot age (based on the year of first plantation) and the percentage of symptomatic plants, although the age range considered was rather small. However, there have been no studies of a possible non-linear effect of age on wood diseases in perennial crops other than vines. Clearly, exploring the underlying mechanisms of wood disease resistance or tolerance according to plant age is a promising research avenue that could improve our understanding of the pathogenesis of such diseases.

By constituting and unifying large spatial and temporal surveillance databases, as in this study, it is possible to visualise and statistically quantify the long-term progression of wood diseases. However, some biases associated with the selection of the plots monitored must be taken into account. In particular, in some wine regions, plots planted with particular cultivars were selected on the basis of their susceptibility to disease rather than as a representative sample of the cultivars in the region. The use of different regional strategies may have decreased the representativeness of the data. This bias would result in a low or high estimated incidence in the region, distorted by the selection of the cultivars monitored. Thus, in such surveillance databases, incidence at regional levels should be interpreted with caution, with particular attention paid to the regional sampling strategy. These potential biases do not call into question the relevance and urgent need for the combination and unification of different regional datasets to develop a national epidemiological surveillance programme for complex diseases, such as esca, in order to help performing additional statistical analyses in both space and time. Pursuing efforts to develop surveillance initiatives and improve both quantity and quality of data is a crucial challenge in the context of global changes. Moreover, climate projections may have far-reaching long-term effects that can only be understood in the light of such long-term surveys.

## Acknowledgement

The authors thank the field observers and farm advisors who collected the data for the National Grapevine Wood Disease Survey and who have continued monitoring at regional level. In particular, we thank the Chambers of Agriculture of Saône-et-Loire, Yonne, Jura, Côte d’Or, Hérault, Rhône, Charente-Maritime and the regional Chamber of Bourgogne-Franche-Comté; the ‘Groupement de Défense contre les Organismes Nuisibles’ (GDON) ‘du libournais’ and ‘des Bordeaux’, the ‘Interprofession des vins du Val de Loire’ (InterLoire), the ‘Comité interprofessionnel du vin de Champagne’ (CIVC), the ‘Bureau interprofessionnel des vins de Bourgogne’ (BIVB); the ‘Institut Français de la Vigne et du Vin’ (IFV); Vitinnov; the ‘Service Régional de l’Alimentation - Direction Régionales de l’Alimentation, de l’Agriculture et de la Forêt’ (SRAL-DRAAF) of Nouvelle-Aquitaine and Auvergne-Rhône-Alpes; the ‘Centre de Recherche Viticole de Corse’ (CRVI); and the ‘Comité National des Interprofessions des Vins’ (CNIV). In addition, we would like to thank the French Epidemiological Plant Health Surveillance Platform (ESV Platform, https://plateforme-esv.fr/) for hosting the database on their information system. We thank Julie Sappa from Alex Edelman & Associates, for the English proofreading of the manuscript. Finally, we would like to thank Thomas Opitz from the “Biostatistiques et Processus Spatiaux” INRAE laboratory for his insight into INLA modelling and Thibaut Fréjaville for preliminary work on the project.

## Funding

This study was supported by the CLIMESCA project (FranceAgriMer grant 22001641) under the framework of the ‘Plan National Dépérissement du Vignoble’.

## Competing interests

The authors declare that they have no competing interests.

## Data availability statement

Part of the research data is available by contacting the corresponding author and a public data visualisation application is available: https://observatoire.plan-deperissement-vigne.fr/adws/app/158a1259-b9ea-11ee-ac62-2997ecd1d68c/

## Notes

### Competing Interest Statement

The authors have declared no competing interest.

### Summary of Updates

This new version was prepared to correct some minor edits.

## References

Aguayo, J., Husson, C., Chancerel, E., Fabreguettes, O., Chandelier, A., Fourrier-Jeandel, C., et al. (2021) Combining permanent aerobiological networks and molecular analyses for large-scale surveillance of forest fungal pathogens: A proof-of-concept. Plant Pathology, 70, 181–194. 10.1111/ppa.13265.

Allen, C.D. (2009) Climate-induced forest dieback: an escalating global phenomenon? 60.

Allen, C.D., Macalady, A.K., Chenchouni, H., Bachelet, D., McDowell, N., Vennetier, M., et al. (2010) A global overview of drought and heat-induced tree mortality reveals emerging climate change risks for forests. Forest Ecology and Management, 259, 660–684. 10.1016/j.foreco.2009.09.001.

Bertsch, C., Larignon, P., Farine, S., Clément, C. & Fontaine, F. (2009) The Spread of Grapevine Trunk Disease. Science, 324, 721–721. 10.1126/science.324_721a.

Bettenfeld, P., Fontaine, F., Trouvelot, S., Fernandez, O. & Courty, P.-E. (2020) Woody Plant Declines. What’s Wrong with the Microbiome? Trends in Plant Science, 25, 381–394. 10.1016/j.tplants.2019.12.024.

Bortolami, G., Gambetta, G.A., Cassan, C., Dayer, S., Farolfi, E., Ferrer, N., et al. (2021) Grapevines under drought do not express esca leaf symptoms. Proceedings of the National Academy of Sciences, 118, e2112825118. 10.1073/pnas.2112825118.

Bruez, E., Baumgartner, K., Bastien, S., Travadon, R., Guérin-Dubrana, L. & Rey, P. (2016) Various fungal communities colonise the functional wood tissues of old grapevines externally free from grapevine trunk disease symptoms. Australian Journal of Grape and Wine Research, 22, 288–295. 10.1111/ajgw.12209.

Bruez, E., Lecomte, P., Grosman, J., Doublet, B., Bertsch, C., Fontaine, F., et al. (2013) Overview of grapevine trunk diseases in France in the 2000s. Phytopathologia Mediterranea, 52, 262–275.

Cailleret, M., Dakos, V., Jansen, S., Robert, E.M.R., Aakala, T., Amoroso, M.M., et al. (2019) Early-Warning Signals of Individual Tree Mortality Based on Annual Radial Growth. Frontiers in Plant Science, 9.

Calzarano, F., Osti, F., Baránek, M. & Di Marco, S. (2018) Rainfall and temperature influence expression of foliar symptoms of grapevine leaf stripe disease (esca complex) in vineyards. Phytopathologia Mediterranea, 57, 488–505.

Carlucci, A., Lops, F., Cibelli, F. & Raimondo, M.L. (2015) Phaeoacremonium species associated with olive wilt and decline in southern Italy. European Journal of Plant Pathology, 141, 717–729. 10.1007/s10658-014-0573-8.

Carlucci, A., Raimondo, M.L., Cibelli, F., Phillips, A.J.L. & Lops, F. (2013) Pleurostomophora richardsiae, Neofusicoccum parvum and Phaeoacremonium aleophilum associated with a decline of olives in southern Italy. Phytopathologia Mediterranea, 52, 517–527.

Carvajal-Yepes, M., Cardwell, K., Nelson, A., Garrett, K.A., Giovani, B., Saunders, D.G.O., et al. (2019) A global surveillance system for crop diseases. Science, 364, 1237–1239. 10.1126/science.aaw1572.

Chacón-Vozmediano, J.L., Gramaje, D., León, M., Armengol, J., Moral, J., Izquierdo-Cañas, P.M., et al. (2021) Cultivar Susceptibility to Natural Infections Caused by Fungal Grapevine Trunk Pathogens in La Mancha Designation of Origin (Spain). Plants, 10, 1171. 10.3390/plants10061171.

Chuine, I., Yiou, P., Viovy, N., Seguin, B., Daux, V. & Ladurie, E.L.R. (2004) Grape ripening as a past climate indicator. Nature, 432, 289–290. 10.1038/432289a.

Ciesla, W.M. & Donaubauer, E. (1994) Decline and Dieback of Trees and Forests: A Global Overview. Food & Agriculture Org.

Cui, J., Li, X., Lu, Z. & Jin, B. (2024) Plant secondary metabolites involved in the stress tolerance of long-lived trees. Tree Physiology, 44, tpae002. 10.1093/treephys/tpae002.

Dewasme, C., Mary, S., Darrieutort, G., Roby, J.-P. & Gambetta, G.A. (2022) Long-Term Esca Monitoring Reveals Disease Impacts on Fruit Yield and Wine Quality. Plant Disease, 106, 3076–3082. 10.1094/PDIS-11-21-2454-RE.

Di Marco, S. & Osti, F. (2008) Foliar Symptom Expression of Wood Decay in Actinidia deliciosa in Relation to Environmental Factors. Plant Disease, 92, 1150–1157. 10.1094/PDIS-92-8-1150.

Dissanayake, A.J., Purahong, W., Wubet, T., Hyde, K.D., Zhang, W., Xu, H., et al. (2018) Direct comparison of culture-dependent and culture-independent molecular approaches reveal the diversity of fungal endophytic communities in stems of grapevine (Vitis vinifera). Fungal Diversity, 90, 85–107. 10.1007/s13225-018-0399-3.

Ficke, A., Gadoury, D.M. & Seem, R.C. (2002) Ontogenic resistance and plant disease management: a case study of grape powdery mildew. Phytopathology, 92, 671–675. 10.1094/PHYTO.2002.92.6.671.

Fischer, M. & Peighami-Ashnaei, S. (2019) Grapevine, esca complex, and environment: the disease triangle. Phytopathologia Mediterranea, 58, 17–37.

Fournier, P., Pellan, L., Barroso-Bergadà, D., Bohan, D.A., Candresse, T., Delmotte, F., et al. (2022) Chapter Two - The functional microbiome of grapevine throughout plant evolutionary history and lifetime. In: Bohan, D.A. and Dumbrell, A. (Eds.) Advances in Ecological Research Functional Microbiomes. Academic Press, pp. 27–99.

Fulchin, E., Verpy, A., Bastiat, C., Bentéjac, S., Coutard, M. & Inchboard, L. (2019) Observatoire des Maladies du Bois sur le vignoble girondin, rapport technique 2021.

Fussler, L., Kobes, N., Bertrand, F., Maumy, M., Grosman, J. & Savary, S. (2008) A Characterization of Grapevine Trunk Diseases in France from Data Generated by the National Grapevine Wood Diseases Survey. Phytopathology®, 98, 571–579. 10.1094/PHYTO-98-5-0571.

Gastou, P., Destrac Irvine, A., Arcens, C., Courchinoux, E., This, P., van Leeuwen, C., & Delmas, C.E.L. (2024) Large gradient of susceptibility to esca disease revealed by long-term monitoring of 46 grapevine cultivars in a common garden vineyard. OENO One, 58(2). 10.20870/oeno-one.2024.58.2.8043

Gramaje, D., Úrbez-Torres, J.R. & Sosnowski, M.R. (2018) Managing Grapevine Trunk Diseases With Respect to Etiology and Epidemiology: Current Strategies and Future Prospects. Plant Disease, 102, 12–39. 10.1094/PDIS-04-17-0512-FE.

Grau, J., Grosse, I. & Keilwagen, J. (2015) PRROC: computing and visualizing precision-recall and receiver operating characteristic curves in R. Bioinformatics, 31, 2595–2597. 10.1093/bioinformatics/btv153.

Grosman, J. & Doublet, B. (2012) Maladies du bois de la vigne: Synthèse des dispositifs d’observation au vignoble, de l’observatoire 2003-2008 au réseau d’épidémiosurveillance actuel. Maladies du bois de la vigne: Synthèse des dispositifs d’observation au vignoble, de l’observatoire 2003-2008 au réseau d’épidémiosurveillance actuel, 31–35.

Gubler, W.D., Rolshausen, P.E., Trouillas, F.P., Úrbez-Torres, J.R., Voegel, T., Leavitt, G.M., et al. (2005) Grapevine trunk diseases in California. Practical Winery and Vineyard, 25.

Guerin-Dubrana, L., Fontaine, F. & Mugnai, L. (2019) Grapevine trunk disease in European and Mediterranean vineyards: occurrence, distribution and associated disease-affecting cultural factors. Phytopathologia Mediterranea, 58, 49–72.

Haffner, V., Enjalric, F., Lardet, L. & Carron, M.P. (1991) Maturation of woody plants: a review of metabolic and genomic aspects. Annales des Sciences Forestières, 48, 615–630. 10.1051/forest:19910601.

Hammond, W.M., Williams, A.P., Abatzoglou, J.T., Adams, H.D., Klein, T., López, R., et al. (2022) Global field observations of tree die-off reveal hotter-drought fingerprint for Earth’s forests. Nature Communications, 13, 1761. 10.1038/s41467-022-29289-2.

Hartmann, H., Moura, C.F., Anderegg, W.R.L., Ruehr, N.K., Salmon, Y., Allen, C.D., et al. (2018) Research frontiers for improving our understanding of drought-induced tree and forest mortality. New Phytologist, 218, 15–28. 10.1111/nph.15048.

Hoo, Z.H., Candlish, J. & Teare, D. (2017) What is an ROC curve? Emergency Medicine Journal, 34, 357–359. 10.1136/emermed-2017-206735.

ISPM 6 (FAO) 1997 Guidelines for surveillance. International Standard for Phytosanitary Measures Publication, 18.

Jurskis, V. (2005) Eucalypt decline in Australia, and a general concept of tree decline and dieback. Forest Ecology and Management, 215, 1–20. 10.1016/j.foreco.2005.04.026.

Kovács, C., Balling, P., Bihari, Z., Nagy, A. & Sándor, E. (2017) Incidence of grapevine trunk diseases is influenced by soil, topology and vineyard age, but not by Diplodia seriata infection rate in the Tokaj Wine Region, Hungary. Phytoparasitica, 45, 21–32. 10.1007/s12600-017-0570-5.

Lecomte, P., Bénétreau, C., Diarra, B., Meziani, Y., Delmas, C., & Fermaud, M. (2024) Logistic modeling of summer expression of esca symptoms in tolerant and susceptible cultivars in Bordeaux vineyards. OENO One, 58.

Lecomte, P., Darrieutort, G., Liminana, J.-M., Comont, G., Muruamendiaraz, A., Legorburu, F.-J., et al. (2012) New Insights into Esca of Grapevine: The Development of Foliar Symptoms and Their Association with Xylem Discoloration. Plant Disease, 96, 924–934. 10.1094/PDIS-09-11-0776-RE.

Lecomte, P., Diarra, B., Carbonneau, A., Rey, P. & Chevrier, C. (2018) Esca of grapevine and training practices in France: results of a 10-year survey. Phytopathologia Mediterranea, 57, 472– 487.

Lecomte, P., Louvet, G., Vacher, B. & Guilbaud, P. (2006) Survival of fungi associated with grapevine decline in pruned wood after composting. Phytopathologia Mediterranea, 45, S127– S130.

Makowski, D., Ben-Shachar, M. & Lüdecke, D. (2019) bayestestR: Describing Effects and their Uncertainty, Existence and Significance within the Bayesian Framework. Journal of Open Source Software, 4, 1541. 10.21105/joss.01541.

Mariette, N., Hotte, H., Chappé, A.-M., Grosdidier, M., Anthoine, G., Sarniguet, C., et al. (2023) Two decades of epidemiological surveillance of the pine wood nematode in France reveal its absence despite suitable conditions for its establishment. Annals of Forest Science, 80, 21. 10.1186/s13595-023-01186-8.

Molyneux, R.J., Mahoney, N., Bayman, P., Wong, R.Y., Meyer, K. & Irelan, N. (2002) Eutypa Dieback in Grapevines: Differential Production of Acetylenic Phenol Metabolites by Strains of Eutypa lata. Journal of Agricultural and Food Chemistry, 50, 1393–1399. 10.1021/jf011215a.

Mondello, V., Songy, A., Battiston, E., Pinto, C., Coppin, C., Trotel-Aziz, P., et al. (2018) Grapevine Trunk Diseases: A Review of Fifteen Years of Trials for Their Control with Chemicals and Biocontrol Agents. Plant Disease, 102, 1189–1217. 10.1094/PDIS-08-17-1181-FE.

Mugnai, L., Graniti, A. & Surico, G. (1999) Esca (Black Measles) and Brown Wood-Streaking: Two Old and Elusive Diseases of Grapevines. Plant Disease, 83, 404–418. 10.1094/PDIS.1999.83.5.404.

Murolo, S. & Romanazzi, G. (2014) Effects of grapevine cultivar, rootstock and clone on esca disease. Australasian Plant Pathology, 43, 215–221. 10.1007/s13313-014-0276-9.

Nutter, F.W., Esker, P.D. & Netto, R.A.C. (2006) Disease Assessment Concepts and the Advancements Made in Improving the Accuracy and Precision of Plant Disease Data. European Journal of Plant Pathology, 115, 95–103. 10.1007/s10658-005-1230-z.

Pandey, P., Ramegowda, V. & Senthil-Kumar, M. (2015) Shared and unique responses of plants to multiple individual stresses and stress combinations: physiological and molecular mechanisms. Frontiers in Plant Science, 6.

Parnell, S., Bosch, F. van den, Gottwald, T. & Gilligan, C.A. (2017) Surveillance to Inform Control of Emerging Plant Diseases: An Epidemiological Perspective. Annual Review of Phytopathology, 55, 591–610. 10.1146/annurev-phyto-080516-035334.

Péros, J.-P., Berger, G. & Jamaux-Despréaux, I. (2008) Symptoms, Wood Lesions and Fungi Associated with Esca in Organic Vineyards in Languedoc-Roussillon (France). Journal of Phytopathology, 156, 297–303. 10.1111/j.1439-0434.2007.01362.x.

Pollastro, S., Faretra, F., Abbatecola, A. & Dongiovanni, C. (2000) Observations on the Fungi Associated with Esca and on Spatial Distribution of Esca-Symptomatic Plants in Apulian (Italy) Vineyards. Phytopathologia Mediterranea, 1000–1005. 10.1400/57845.

Reisenzein, H., Nieder, G. & Berger, N. (2000) Esca in Austria. Esca in Austria, 1000–1009. 10.1400/57806.

Romanazzi, G., Murolo, S., Pizzichini, L. & Nardi, S. (2009) Esca in young and mature vineyards, and molecular diagnosis of the associated fungi. European Journal of Plant Pathology, 125, 277–290. 10.1007/s10658-009-9481-8.

Rue, H., Martino, S. & Chopin, N. (2009) Approximate Bayesian Inference for Latent Gaussian models by using Integrated Nested Laplace Approximations. Journal of the Royal Statistical Society Series B: Statistical Methodology, 71, 319–392. 10.1111/j.1467-9868.2008.00700.x.

Rue, H., Riebler, A., Sørbye, S.H., Illian, J.B., Simpson, D.P. & Lindgren, F.K. (2017) Bayesian Computing with INLA: A Review. Annual Review of Statistics and Its Application, 4, 395–421. 10.1146/annurev-statistics-060116-054045.

Serra, S., Ligios, V., Schianchi, N., Prota, V.A. & Scanu, B. (2018) Expression of grapevine leaf stripe disease foliar symptoms in four cultivars in relation to grapevine phenology and climatic conditions. Phytopathologia Mediterranea, 57, 557–568.

Sosnowski, M.R., Shtienberg, D., Creaser, M.L., Wicks, T.J., Lardner, R. & Scott, E.S. (2007) The Influence of Climate on Foliar Symptoms of Eutypa Dieback in Grapevines. Phytopathology®, 97, 1284–1289. 10.1094/PHYTO-97-10-1284.

Surico, G., Mugnai, L., Braccini, P. & Marchi, G. (2000) Epidemiology of Esca in Some Vineyards in Tuscany (Italy). Phytopathologia Mediterranea, 1000–1016. 10.1400/57844.

Úrbez-Torres, J.R., Adams, P., Kamas, J. & Gubler, W.D. (2009) Identification, Incidence, and Pathogenicity of Fungal Species Associated with Grapevine Dieback in Texas. American Journal of Enology and Viticulture, 60, 497–507. 10.5344/ajev.2009.60.4.497.

Valencia, A.L., Saavedra-Torrico, J., Rosales, I.M., Mártiz, J., Retamales, A., Link, A., et al. (2022) Unveiling the Predisposing Factors for the Development of Branch Canker and Dieback in Avocado: A Case of Study in Chilean Orchards. Horticulturae, 8, 1121. 10.3390/horticulturae8121121.

Zuur, A.F., Ieno, E.N. & Saveliev, A.A. (2017) Using GLM and GLMM. Newburgh, United Kingdom: Highland Statistics Ltd.

