## Supplementary materials for "Exploring the role of cultivar, year and plot age in the incidence of esca and Eutypa dieback: insights from 20 years of regional surveys in France"

### Supporting Information

Table S1: Number and percentage of plots monitored for esca and Eutypa dieback leaf symptoms, by French wine region and period. The locations of the French wine regions are shown on Figure 1.

| French wine regions | Monitoring years | Number of plots monitored | Percentage of plots monitored for esca | Percentage of plots monitored for Eutypa dieback |
|---|---|---|---|---|
| Alsace Lorraine | 2003-2022 | 131 | 100 | 70 |
| Bordelais | 2003-2022 | 381 | 100 | 67 |
| Bourgogne | 2003-2022 | 174 | 100 | 61 |
| Champagne | 2003-2022 | 586 | 100 | 0 |
| Charentes | 2003-2022 | 72 | 100 | 89 |
| Corse | 2003-2021 | 54 | 100 | 0 |
| Jura & Savoie | 2004-2022 | 85 | 100 | 100 |
| Languedoc | 2003-2022 | 55 | 100 | 71 |
| Provence | 2003-2012 | 70 | 100 | 34 |
| Sud Ouest | 2003-2007 | 127 | 100 | 2 |
| Val de Loire | 2003-2008 / 2012-2022 | 250 | 100 | 100 |
| Vallée du Rhône | 2003-2019 | 95 | 100 | 53 |
| Vendée | 2003-2008 | 2 | 100 | 100 |

Table S2: Mean number of vines monitored per plot (± standard deviation, SD) and number of plots monitored for each cultivar

| Cultivar | Mean ± SD | Number of plots monitored |
|---|---|---|
| Alphonse Lavallée | 300 ± 0 | 1 |
| Cabernet Franc | 452 ± 303 | 115 |
| Cabernet Sauvignon | 1176 ± 1648 | 143 |
| Carignan | 300 ± 2 | 36 |
| Chardonnay | 351 ± 257 | 344 |
| Chasselas | 299 ± 7 | 25 |
| Chenin | 276 ± 55 | 58 |
| Cinsault | 386 ± 304 | 25 |
| Colombard | 309 ± 39 | 24 |
| Fer Servadou | 393 ± 202 | 27 |
| Gamay | 298 ± 9 | 45 |
| Gewurztraminer | 295 ± 18 | 42 |
| Grenache | 300 ± 0 | 32 |
| Italia | 300 ± 0 | 1 |
| Malbec | 309 ± 59 | 27 |
| Melon | 300 ± 2 | 53 |
| Merlot | 572 ± 188 | 87 |
| Meunier | 330 ± 43 | 1877 |
| Mourvèdre | 355 ± 194 | 9 |
| Muscat de Hambourg | 300 ± 0 | 15 |
| Muscat Petits Grains | 300 ± 2 | 46 |
| Négrette | 300 ± 0 | 25 |
| Niellucciu | 207 ± 11 | 16 |

| | | |
|---|---|---|
| Pinot Auxerrois | 293 ± 21 | 45 |
| Pinot Noir | 392 ± 360 | 193 |
| Poulsard | 300 ± 0 | 25 |
| Riesling | 295 ± 30 | 43 |
| Sauvignon Blanc | 469 ± 339 | 155 |
| Sauvignon Gris | 543 ± 11 | 1 |
| Savagnin | 298 ± 13 | 30 |
| Sciaccarellu | 315 ± 182 | 19 |
| Semillon | 633 ± 143 | 46 |
| Syrah | 335 ± 197 | 24 |
| Trousseau | 298 ± 11 | 30 |
| Ugni Blanc | 385 ± 179 | 72 |
| Vermentinu | 211 ± 8 | 19 |

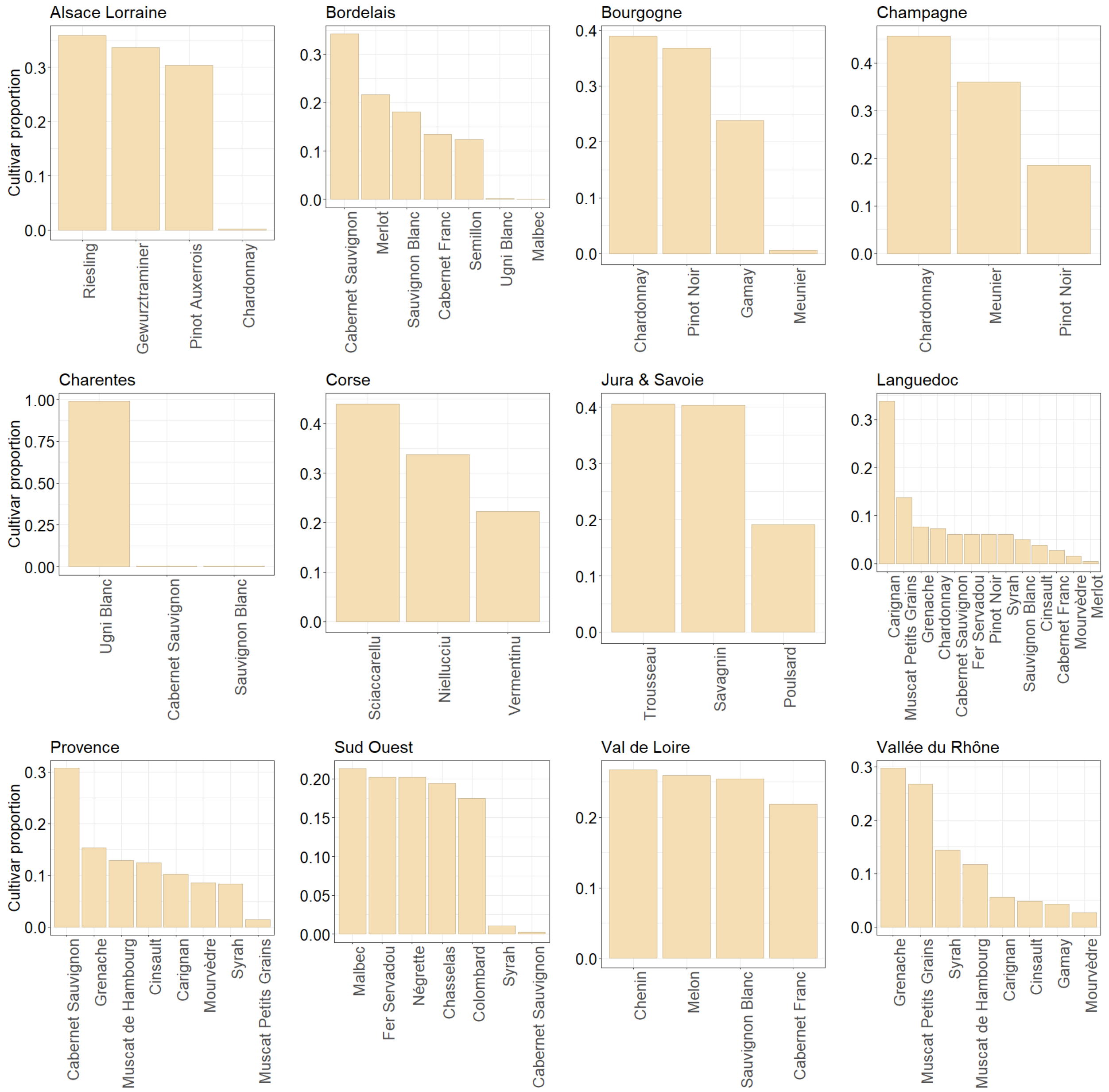


Figure S1: The proportions of the cultivars monitored by the French wine region, from 2003 to 2022. Note that Bourgogne, Vallée du Rhône, Corse, Sud Ouest and Bordelais regions are internationally known as Burgundy, Rhone Valley, Corsica, South West, and Bordeaux, respectively.

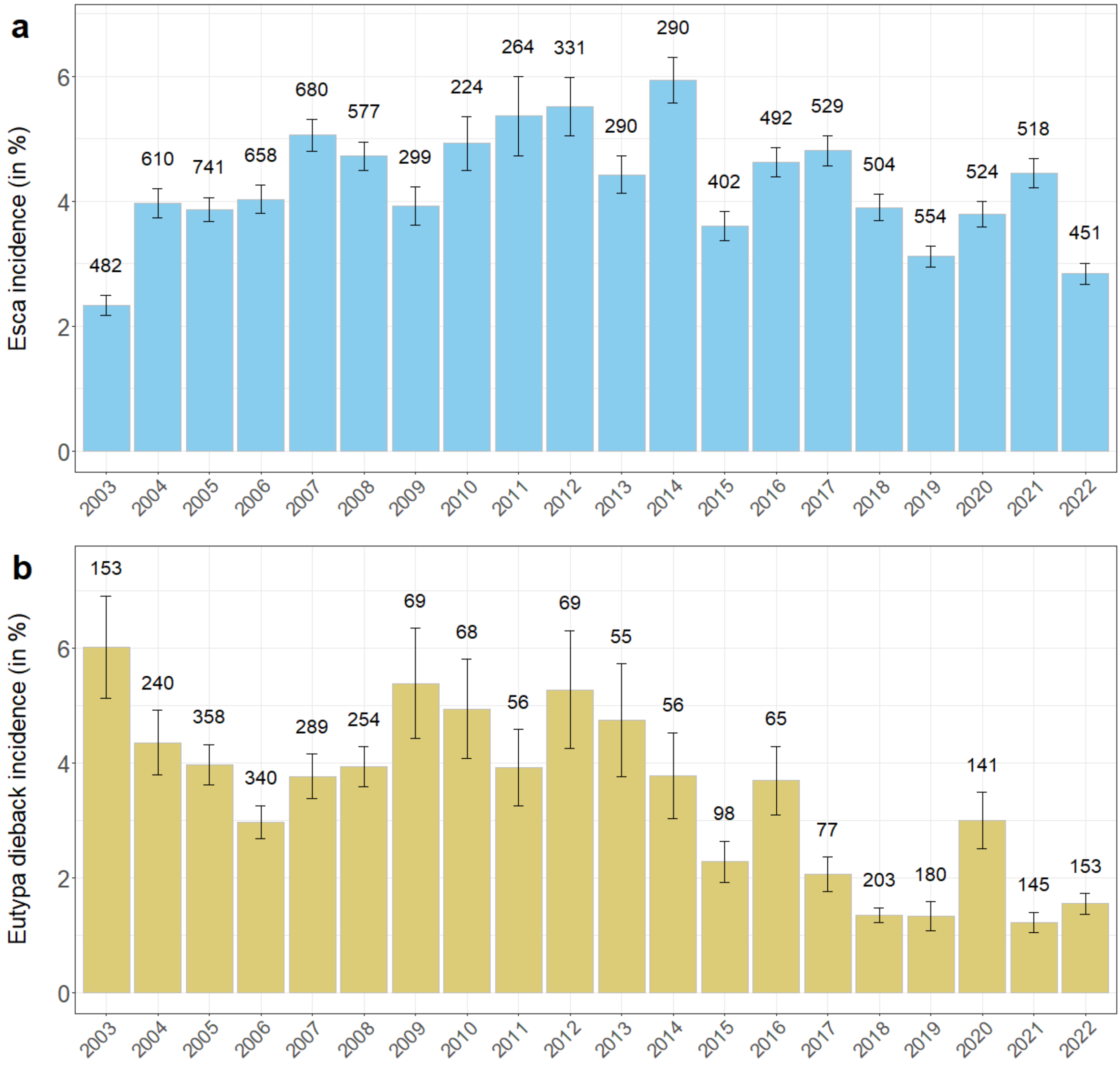


Figure S2: Mean incidence of (a) esca (blue bars) and (b) Eutypa dieback (mustard bars) in France over the period 2003-2022, excluding plots on which no grapevine trunk diseases were observed (prevalence > 0). The incidence is the percentage of symptomatic plants observed per plot and per year. The number of plots monitored per year and per disease are indicated above the bars. The error bars represent the standard error of the mean.

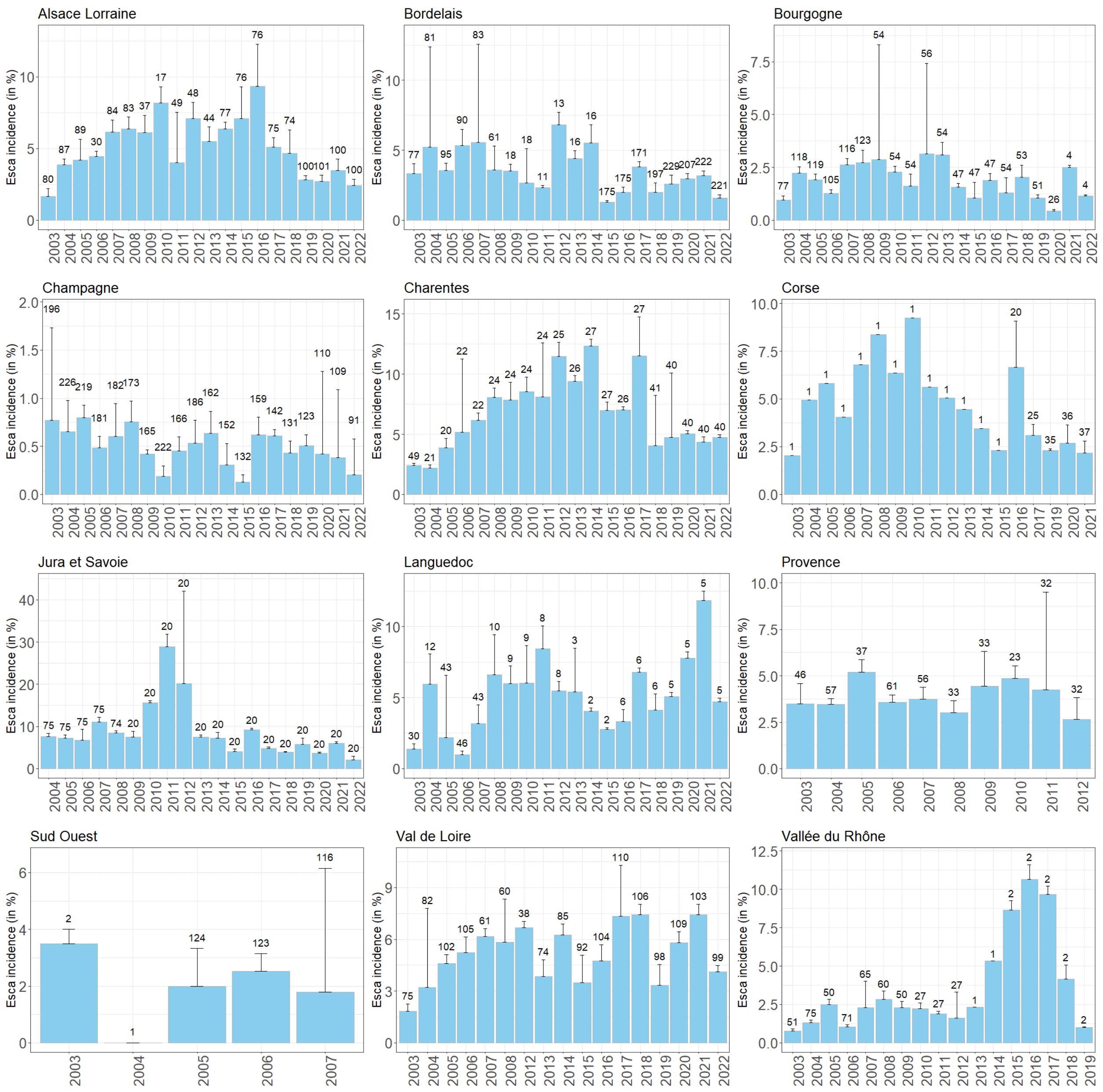


Figure S3: The mean incidence of esca for each wine region monitored in France for the 2003-2022 period. The number of plots monitored is indicated above each bar. The error bars represent the standard errors of the mean, with only positive errors shown. Note that the scale of the *y*-axis differs between regions.

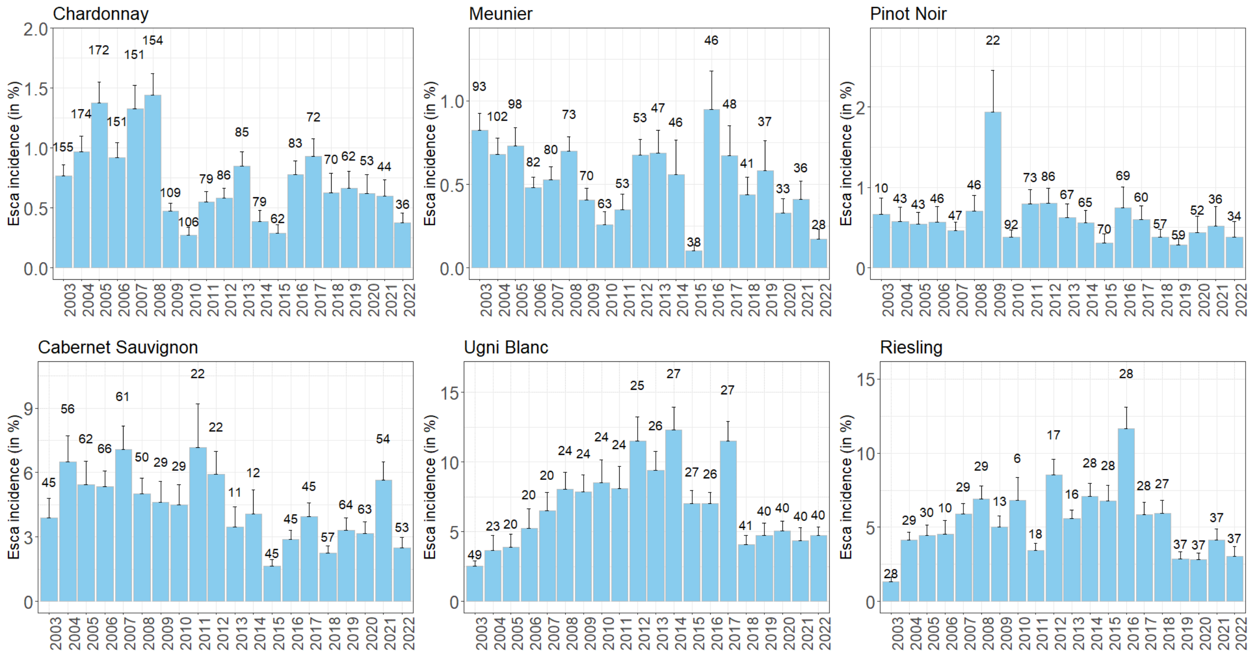


Figure S4: Mean incidence of esca between 2003 and 2022 for the six cultivars with the largest numbers of plots monitored in the whole database. The number of plots monitored is indicated above each bar. Error bars represent the standard error of the mean and only the positive error is shown. Note that the scale of the *y*-axis differs between cultivars.

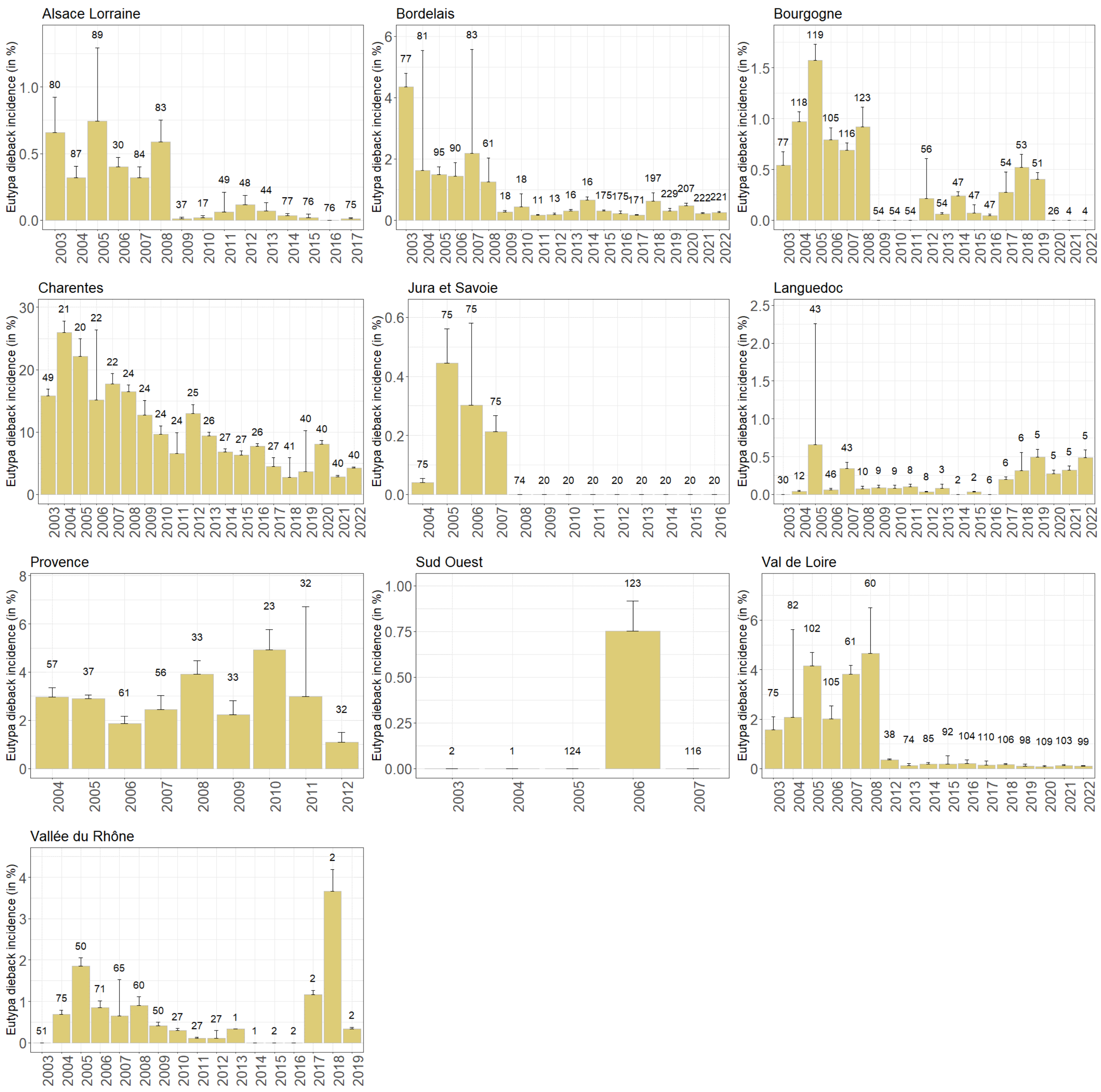


Figure S5: Mean incidence of Eutypa dieback in French wine regions monitored over the 2003-2022 period. The number of plots monitored is indicated above each bar. The error bars represent the standard error of the mean, with only the positive error shown. Note that the scale of the *y*-axis differs between regions.

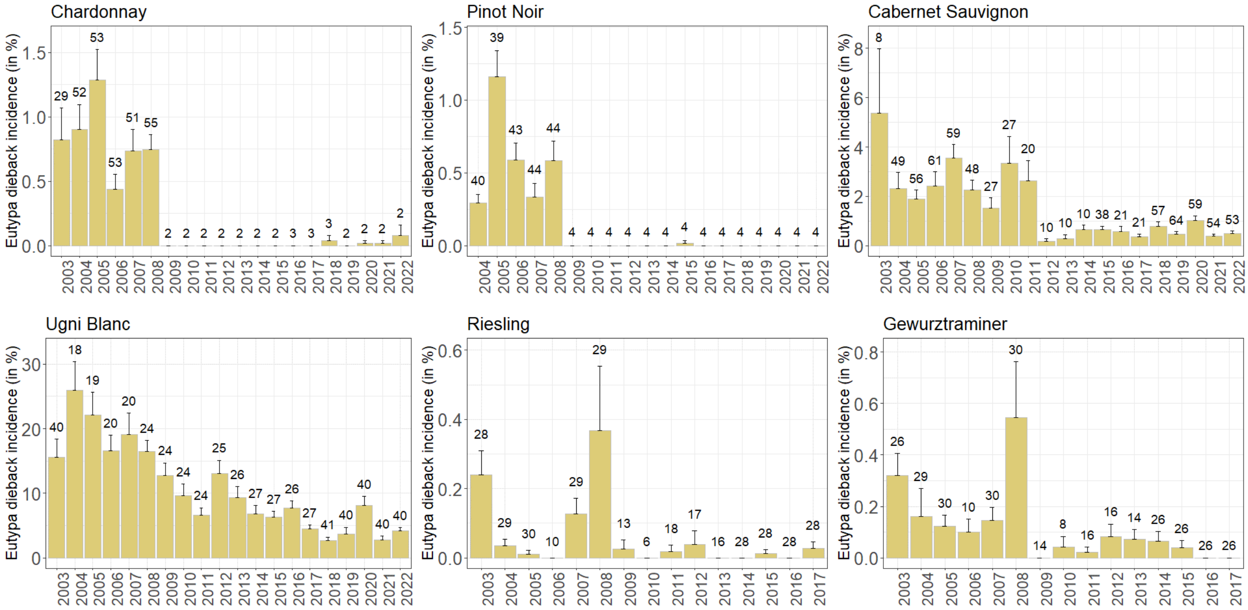


Figure S6: Mean incidence of Eutypa dieback in the 6 cultivars for which the largest numbers of plots were monitored for the 2003-2022 period. The number of plots monitored is indicated above each bar. The error bars represent the standard error of the mean, with only the positive error shown. Note that the scale of the *y*-axis differs between cultivars.
